# p53 and YAP/TAZ-TEAD activities determine metaplastic heterogeneity in pancreatic cancer

**DOI:** 10.64898/2025.12.11.693520

**Authors:** Cole P. Martin, Scott Bang, Joseph R. Gomes, William B. Sullivan, Yusha Liu, Matthew D. Smith, A. Cole Edwards, Channing J. Der, John P. Morris

## Abstract

**BACKGROUND and AIMS:** Pancreatic ductal adenocarcinoma (PDAC) initiation and progression is characterized by lineage plasticity across a continuum of metaplastic and malignant cell fates. This molecular heterogeneity influences disease progression and therapeutic response, yet the genetic and molecular interactions underlying cellular plasticity during PDAC progression remain poorly understood. Here we studied how p53, the most frequently inactivated tumor suppressor in PDAC, interacts with the YAP/TAZ-TEAD transcriptional effector arm of the Hippo pathway to determine cell fate and neoplastic heterogeneity in PDAC development.

**METHODS:** We characterized differentiation states in mouse PDAC and pre-malignant cells in response to p53 activation or inactivation, respectively, and upon genetic, pharmacological, and physiological modulation of YAP/TAZ-TEAD activity. Differentiation was analyzed *in vivo* in an orthotopic PDAC model permitting independent, inducible control of p53 and TEAD function. We analyzed the effect of p53 and YAP/TAZ-TEAD activity on cell fate in mouse and human PDAC and analyzed markers of clinically relevant molecular subtypes in human PDAC cell lines treated with small molecule TEAD inhibitors.

**RESULTS:** Re-engaging wildtype p53 function in Kras^Mut^ pancreatic cancer cells increases TEAD activity through accumulation of its co-activator TAZ. Remarkably, increased YAP/TAZ-TEAD activity broadly suppresses the expression of overlapping markers of gastric pit-like (GPL) differentiation and the classical PDAC subtype that are promoted by p53. Accordingly, p53 and YAP/TAZ-TEAD activity inversely correlate with GPL and classical differentiation during PDAC development and modulation of both pathways determines expression of GPL and classical markers *in vitro* and *in vivo*. GPL differentiation is acutely sensitive to YAP/TAZ-TEAD activity including oncogenic perturbations of the Hippo pathway, actin dynamics, cell-to-cell contact, and adaptative YAP/TAZ accumulation in response to Ras/MAPK pathway inhibition. Functional repression of gastric-classical differentiation by TEAD persists in human PDAC where potent TEAD inhibition increases expression of GPL and classical markers.

**CONCLUSIONS:** We find that p53 and YAP/TAZ-TEAD activities determine gastric plasticity during PDAC development highlighting the role that key cancer drivers and therapeutically targetable signaling pathways play in active maintenance of cellular identity during pancreatic cancer development.

## INTRODUCTION

Pancreatic ductal adenocarcinoma (PDAC) remains a significant clinical challenge with a 13% 5-year survival rate and a lack of consistently effective therapies^1^. Poor prognosis and therapeutic resistance in the disease is associated with pervasive transcriptional and molecular heterogeneity^2,3^. Advanced disease presents on a plastic spectrum of convertible “classical” and “basal” transcriptional identities, associated with distinct patterns of molecular and histopathological differentiation^4–10^. The classical subtype is linked with epithelial and gastric differentiation genes while basal stratifying genes reflect squamous differentiation and epithelial to mesenchymal transition (EMT)^4–7,11^. Similarly, heterogeneous differentiation states are observed in the earliest stages of PDAC initiation. Histological and single cell analyses have revealed that metaplastic, pre-malignant cells represent a spectrum of duct-like populations that express distinct patterns of endoderm differentiation genes, including markers associated with stomach differentiation such as chief and gastic pit markers^11–13^. As heterogeneous, pre-malignant states may represent different paths to malignancy^11,14–17^ and the classical-basal axis is associated with both PDAC progression and therapeutic response^3–5,8,18^, understanding the molecular triggers underlying plasticity between heterogeneous cell fates during PDAC development has implications for diagnostic and therapeutic strategies.

The PDAC cancer driver gene landscape is well characterized but cannot fully explain the transcriptional heterogeneity observed in advanced disease or pre-malignant metaplasia^19^. PDAC is characterized by a largely uniform set of recurrent mutations, including nearly universal oncogenic mutations in the proto-oncogene Kras, and frequent inactivating mutations in the tumor suppressors TP53, SMAD4, and CDKN2A^20^. While distinct patterns of driver genes often define transcriptional heterogeneity in cancers^21,22^, classical or basal identity is not an exclusive feature of specific combinations of Kras and tumor suppressor mutations in PDAC, but rather, the classical-basal continuum is observed across driver configurations^19^. Furthermore, pre-malignant heterogeneity along the ductal-gastric axis is associated with initiating Kras mutations (Kras^Mut^) in human disease^23–25^ and can be potently triggered by Kras^Mut^ absent additional mutations in mouse models^12,13,26,27^. Therefore, while PDAC driver mutations enable the acquisition of aberrant cell fates during PDAC development, neoplastic heterogeneity is ultimately determined by the activity of cell signaling pathways and transcriptional regulators that are inappropriately activated during tumorigenesis. How PDAC driver genes interact with these determinants of heterogeneity remains poorly understood.

p53 is the most frequently mutated tumor suppressor in PDAC, co-occurring with Kras^Mut^ in ∼75% of cases^20^. p53 inactivation is strongly associated with the progression from well-differentiated pre-malignant cells to frank, invasive cancer and the loss of epithelial and endoderm differentiation characteristic of cancer precursors^25,28^. This role is conserved in mouse models initiated by Kras^Mut^ and p53 inactivation that recapitulate seminal features of pre-malignant to malignant progression^29,30^. We have previously demonstrated that p53 controls cellular plasticity in PDAC as restoration of wildtype p53 in advanced murine disease re-establishes epithelial differentiation and elements of metaplastic, pre-malignant gene expression^31^. Here, we find that p53 determines metaplastic gene expression via interactions with the key transcriptional effectors of the HIPPO pathway, YAP/TAZ and TEAD^32^. We find that p53 promotes the expression of gastric pit-like differentiation programs overlapping with those observed in the classical PDAC subtype that are repressed by YAP/TAZ-TEAD activity. Modulating p53 and TEAD activity via genetic and physiological regulators of YAP and TAZ thus determines expression of gastric and classical differentiation genes both in vitro and in vivo. YAP/TAZ-TEAD activity is inversely correlated with gastric and classical differentiation in PDAC, and the relationship between signaling and differentiation is dynamic. Markers of gastric and classical differentiation are upregulated in response to pharmacological TEAD inhibition and suppressed by adaptive accumulation of YAP/TAZ in response to inhibition of the KRAS/MAPK pathway. Thus, our work reveals the ability of a key PDAC tumor suppressor and a targetable transcriptional regulator that is co-opted during PDAC progression to determine metaplastic heterogeneity and cellular plasticity during disease progression and therapeutic response.

## RESULTS

### p53 and YAP/TAZ activities compete to determine expression of gastric pit-like and classical subtype differentiation signatures in pancreatic cancer cells

p53 status is associated with changes in cell fate during PDAC progression where its inactivation leads to the progression of well-differentiated, pre-malignant cells to transcriptionally distinct invasive disease^30,33^. To study the role that p53 plays in defining cell fate during PDAC development, we previously developed the “KP^sh”^ model. Here, advanced disease is initiated by pancreas specific expression of oncogenic Kras^Mut^ and a doxycycline (dox) inducible shRNA targeting p53 (*KP^sh^*: *Pdx1-Cre; LSL-Kras^G12D^; TRE-shp53-GFP; R26-LSL-rtTA-mKate2*)^31^(Figure 1A). Withdrawal of dox from primary KP^sh^ PDAC cells results in robust accumulation of endogenously expressed, transcriptionally active p53 (Figure 1B) that triggers increased expression of markers of epithelial and pre-malignant differentiation^31^, suggesting an active role for p53 in determining cellular plasticity during PDAC progression.

**Figure 1.**
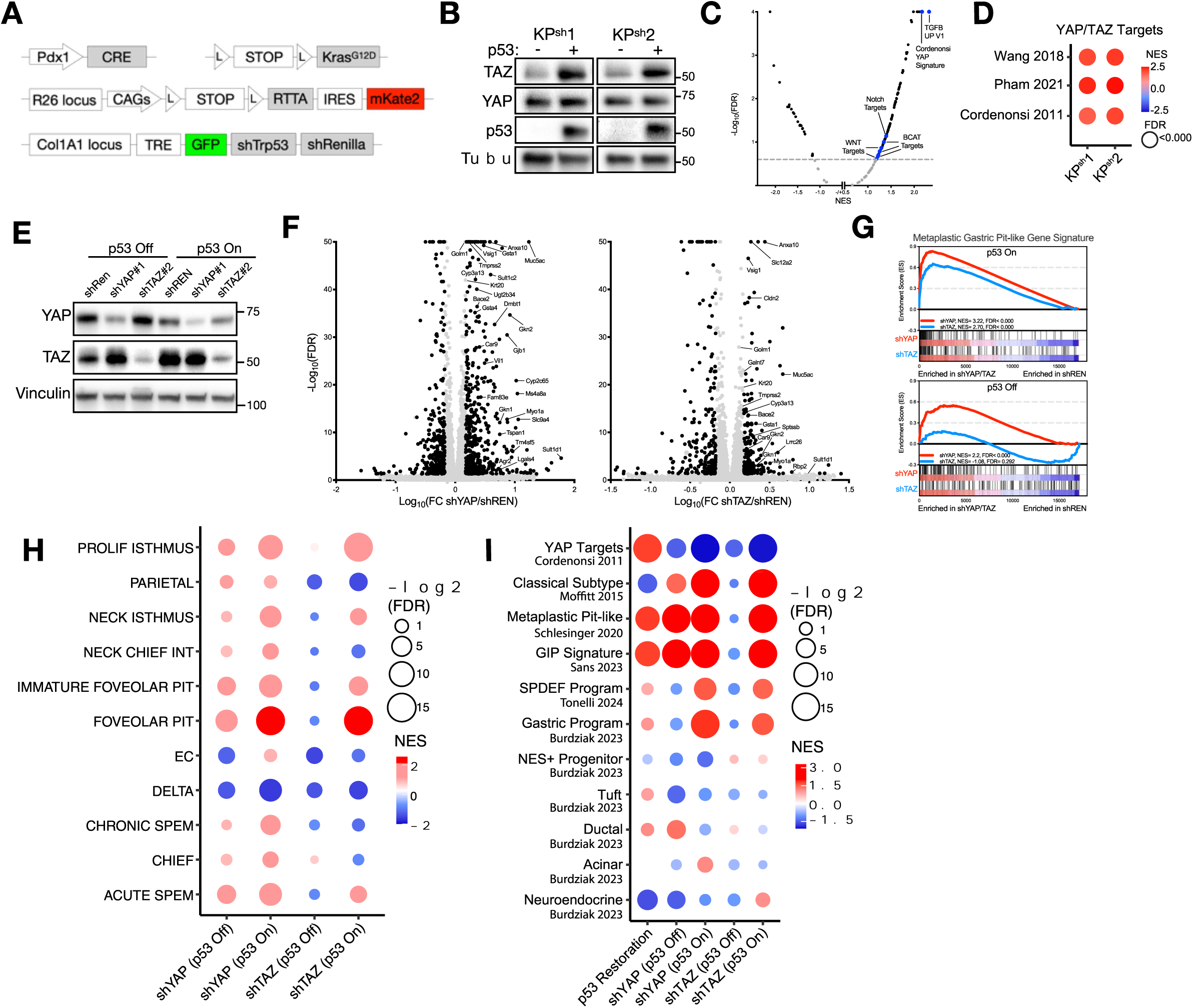
p53 and YAP/TAZ activities compete to determine gastric and classical subtype signature expression in pancreatic cancer cells. **(A)** Schematic of the KP^sh^ model alleles. **(B)** Western blots of KP^sh^ cells. **(C)** GSEA of all oncogenic signatures (MSigDB C6) from RNA-seq in KP^sh^2 cells. **(D)** GSEA of three YAP/TAZ target gene signatures upon p53 restoration. **(E)** Western blots of KP^sh^1 expressing shRNAs. **(F-I)** RNA-seq of KP^sh^1 cells expressing shRNAs targeting Yap1 (YAP), Wwtr1 (TAZ), or Renilla luciferase (REN, control). **(F)** Volcano plots with p53 on. **(G-I)** GSEA of published **(G)** metaplastic gastric pit-like genes **(H)** stomach cell derived gene sets, and **(I)** other relevant gene sets associated with pancreatic metaplastic heterogeneity.

Cell fate during PDAC development is shaped by transcription factors involved in pancreatic development and regeneration that are inappropriately activated during disease initiation and progression^,34^. Remarkably, gene set enrichment analysis (GSEA) of RNA-SEQ profiling following sustained p53 restoration in KP^sh^ cells revealed enrichment of target genes of several transcriptional mediators that have been implicated in neoplastic cell fate. This included targets of the TGF-β^35,36^, Notch^37,38^, and Wnt/β-Catenin pathways^39,40^, as well as those associated with activity of YAP and TAZ, transcriptional co-activators of the TEAD family of transcription factors whose levels are mediated by the HIPPO pathway^41,42^ (Figure 1C). p53 restoration increased expression of multiple YAP/TAZ target signatures (Figure 1D) and resulted in increased levels of TAZ (Figure 1B) in KP^sh^ cells. Given the important role that YAP and TAZ plays in PDAC initiation^41^, maintenance^42^, and therapeutic response^43^ we set out to determine the effect of increased YAP/TAZ transcriptional activity on p53 dependent gene expression.

To address this question, we performed RNA-seq in p53 silenced and restored KP^sh^ cells expressing constitutive shRNAs that effectively knocked down YAP or TAZ (Figure 1E). YAP or TAZ knockdown dramatically altered gene expression upon p53 restoration (Figure 1F), including the upregulation of hundreds of genes, suggesting a role for YAP and TAZ in antagonizing gene expression promoted by p53 activity. Remarkably, these derepressed genes were dominated by markers of gastric pit-like (GPL) differentiation, a heterogeneous, metaplastic identity enabled by Kras^Mut^ during pre-malignant development^12,13,27^ that corresponds with both human and mouse PanINs^12,25^ and displays transcriptional overlap with the classical PDAC transcriptional subtype^12^. YAP or TAZ silencing led to upregulation of 71 out of 155 genes (45.8%) comprising a GPL signature defined from single cell expression analysis of Kras^Mut^ pre-malignant epithelium^12^ (Fold change > 1.5, FDR<0.05), indicating a regulatory relationship where p53 and YAP/TAZ activity compete to determine GPL differentiation. GSEA further revealed that this GPL signature was weakly enriched by YAP or TAZ knockdown where p53 remained inactive but was strongly enriched by silencing of both genes in the setting of p53 restoration (Figure 1G). Competition between p53 and YAP/TAZ activity to promote metaplastic gastric cell type expression extended to gene signatures which define other heterogeneous identities in the adult stomach^13,44^, with signatures of mucus producing foveolar pit cells reflecting the highest degree of enrichment upon p53 restoration in the setting of YAP or TAZ knockdown (Figure 1H). However, enteroendocrine differentiation associated with hormone secreting cells (e.g. EC and delta cells) displayed no p53 or YAP/TAZ mediated enrichment (Figure 1H). To better understand the extent of p53-YAP/TAZ competition on neoplastic cell fate, we extended our analysis to metaplastic identities identified in pre-malignant epithelium driven by Kras^mut^. While progenitor, tuft, ductal, and endocrine signatures were not regulated by p53-YAP/TAZ competition, gastric signatures identified from multiple single cell expression studies of Kras^mut^ driven pre-malignant metaplasia, as well as the classical subtype of human PDAC, showed a pronounced upregulation in the setting of YAP/TAZ knockdown, particularly upon p53 restoration (Figure 1I). Taken together, these data suggest that p53 activity preferentially promotes gastric differentiation characteristic of Kras^Mut^ pre-malignant cells and the classical subtype of human PDAC that is antagonized by the action of YAP and TAZ.

### YAP/TAZ activity constrains gastric differentiation promoted by p53

We next validated the ability of p53 and YAP/TAZ activities to control gastric-like differentiation in pancreatic cancer cells. Consistent with expression profiling by RNA-seq, YAP and TAZ knockdown in two independently derived KP^sh^ cell lines resulted in striking increases of GKN2 and MUC5AC protein levels, two markers of GPL differentiation (Figure 2A). Additionally, gene expression of GPL markers Gkn1, Gkn2, Muc5ac, and Tff1 were markedly increased in the setting of YAP and TAZ knockdown following p53 restoration, while levels were only modestly increased by p53 restoration alone (Figure 2B, Supplemental Figure 1A). Of note, this effect depended closely on cell line specific YAP and TAZ levels. Depletion of both YAP and TAZ in KP^sh^2 cells produced equivalent gastric gene upregulation, while TAZ depletion in KP^sh^1 cells resulted in a more modest effect likely owing to the lower TAZ:YAP ratio in this cell line (Figure 2A,B). Increased GPL marker expression was similarly observed in a p53 dependent fashion in KP^sh^ cell lines expressing sgRNAs targeting either YAP or TAZ (Supplemental Figure 1A-D). Accordingly, immunofluorescence and flow cytometry analysis of the GPL marker MUC5AC revealed an increased frequency of marker positive cells in response to p53 restoration that was dramatically potentiated by Yap or Taz knockdown (Figure 2C,D; Supplemental Figure 1E,F).

**Figure 2.**
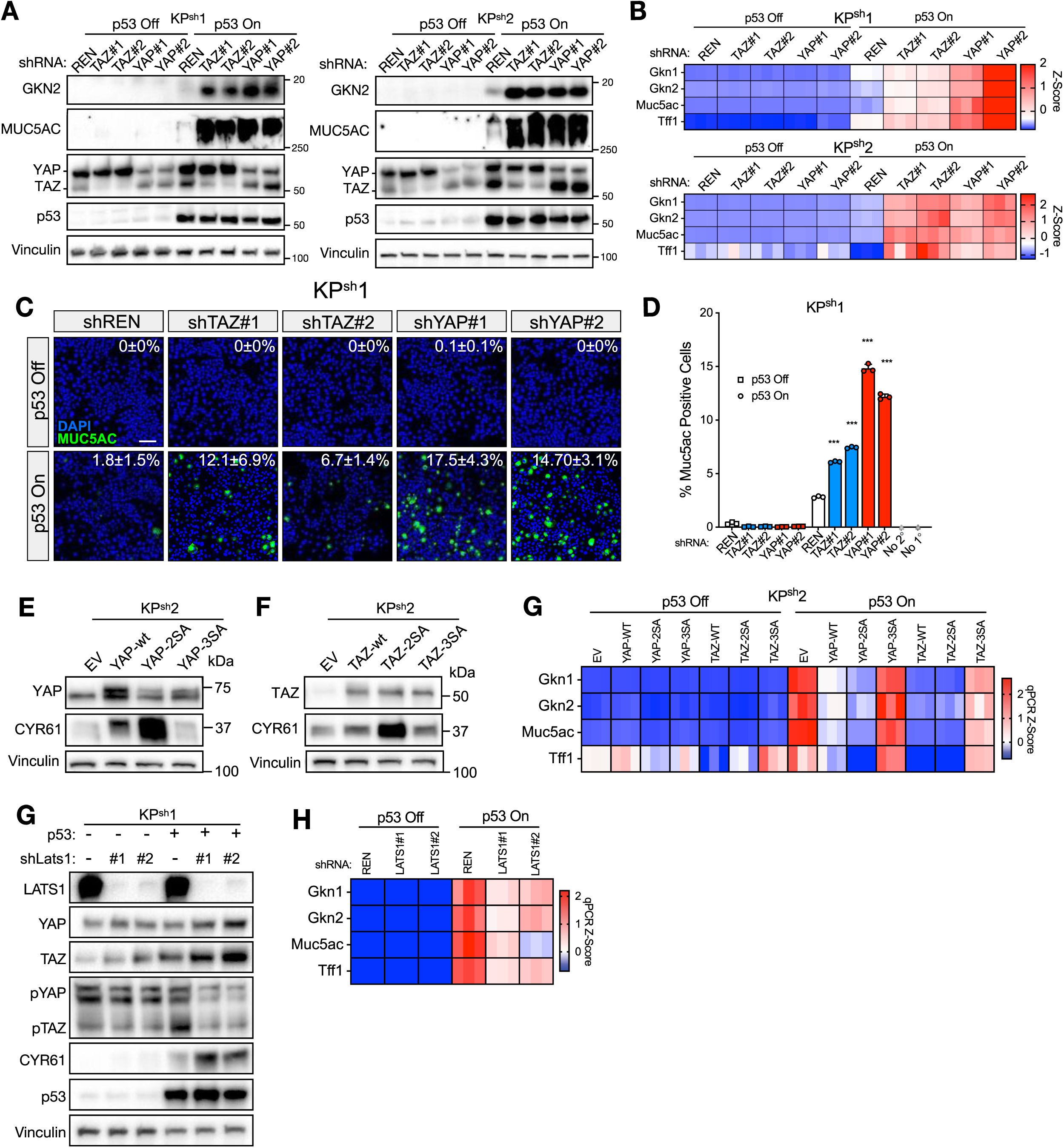
YAP/TAZ activity constrains gastric differentiation promoted by p53. **(A)** Western blots of YAP/TAZ shRNAs and relevant gastric genes. **(B)** RT-qPCRs for gastric genes, normalized to Gusb, converted to Z-scores, and plotted in heatmaps. **(C)** IF microscopy of MUC5AC and DAPI. Upper right corner of images displays percentage MUC5AC positive cells with associated standard deviation across an average of n=5.5 fields representing an average of n=1367 cells per condition. **(D)** Flow cytometry depicting percent MUC5AC positive KP^sh^1 cells from three independent wells. **(E-G)** KP^sh^2 overexpressing indicated alleles. **(E,F)** Representative western blots showing **(E)** YAP and **(F)** TAZ overexpression. **(G)** Heatmaps of RT-qPCRs showing Z-scored expression. **(H,I)** KP^sh^1 cells expressing two shRNAs targeting Lats1. **(H)** Western blots demonstrating LATS1 knockdown. **(I)** Heatmap of Z-scored qRT-qPCR expression.. Scale bar in (C) = 100 µm.

Increased levels of YAP and TAZ activity via genome amplification^45^ or inactivation of their negative regulators^46^ can perturb pancreatic cell fate and drive poorly differentiated, squamous differentiation phenotypes in PDAC^47^. Therefore, we asked if these gain of function effects on YAP or TAZ could repress GPL differentiation triggered by p53 activity. YAP/TAZ levels are controlled by the LATS1/2 kinases, which phosphorylate YAP and TAZ at several serine residues to induce cytoplasmic sequestration and degradation^48^. Thus, we overexpressed either wildtype human YAP and TAZ, hyperactive alleles harboring serine to alanine mutations at the key phosphorylation sites YAP-S127A-S397A (YAP-2SA) and TAZ-S89A-S311A (TAZ-2SA), or inactive forms YAP-S127A-S397A-S94A (YAP-3SA) and TAZ-S89A-S311A-S51A (TAZ-3SA) that only weakly interact with TEAD transcription factors and studied their effects on gastric gene expression in response to p53 restoration (Figure 2E,F, Supplemental Figure 1G,H). Overexpression of both wildtype and hyperactive YAP/TAZ-2SA in KP^sh^ cells blocked Gkn1, Gkn2, Muc5ac, and Tff1 upregulation by p53, which was partially rescued by the TEAD binding deficient 3SA mutant (Figure 2G, Supplemental Figure 1I). Importantly, inhibition of GPL markers directly correlated with induction of the YAP/TAZ transcriptional target Cyr61 (Figure 2E,F, Supplemental Figure 1G,H). This is consistent with the transcriptional consequences of YAP/TAZ interacting with p53 to determine GPL differentiation.

Increasing YAP/TAZ levels by depleting the LATS1 tumor suppressor was also sufficient to repress GPL gene expression in response to p53 activation. shRNA knockdown of Lats1 in KP^sh^ cells resulted in decreased TAZ phosphorylation and increased TAZ levels with more modest effects on YAP phosphorylation and accumulation that correlated directly with expression of CYR61 (Figure 2H). Accordingly, Lats1 knockdown blunted upregulation of Gkn1, Gkn2, Muc5ac, and Tff1 in response to p53 restoration (Figure 2I). Taken together, we demonstrate that YAP/TAZ activity plays a critical role in suppressing GPL differentiation promoted by p53 activity in pancreatic cancer cells.

### YAP/TAZ act through TEAD to suppress p53 triggered gastric differentiation

YAP/TAZ lack DNA binding domains but rather act as transcriptional co-activators of several transcription factors (TFs), including the TEAD(1-4) TF family^49^. YAP/TAZ dependent TEAD activity has been implicated in the specification and maintenance of both Kras^Mut^ pre-malignant metaplasia^41^ and PDAC^42^. Thus, we used pharmacological and dominant negative approaches to test if TEAD activity connects YAP/TAZ to suppression of gastric differentiation promoted by p53. First, we gauged GPL marker expression in KP^sh^ cells treated with two mechanistically distinct small molecule inhibitors of TEAD, VT104 which blocks TEAD-auto-palmitoylation necessary for its activation^50^ and IAG933 which interferes with YAP/TAZ binding to TEAD^51^. Both VT104 and IAG933 downregulated the well-characterized YAP/TAZ-TEAD target CYR61 in KP^sh^ cells when p53 remained silenced and upon p53 restoration (Figure 3A). Demonstrating that YAP and TAZ activity function through TEAD to suppress GPL differentiation, both TEAD inhibitors resulted in a robust upregulation of GKN2 and MUC5AC protein following p53 restoration (Figure 3A). Furthermore, RT-qPCR revealed dose-dependent upregulation of Gkn1, Gkn2, Muc5ac, and Tff1 in response to IAG933 and VT104 upon p53 restoration to levels inaccessible in cells where p53 remained silenced (Figure 3B, Supplementary Figure 2A). Increased GPL marker expression in response to TEAD inhibition corresponded with a similar pattern of emergence of GKN2 and MUC5AC positive subpopulations as observed with YAP and TAZ silencing. Here, p53 restoration or TEAD inhibition alone resulted in rare MUC5AC and GKN2 positivity, with up to ∼30-50% of cells acquiring GPL marker expression (e.g. MUC5AC) in response to TEAD inhibition upon p53 activation (Figure 3C,D; Supplemental Figure 2B,C).

**Figure 3.**
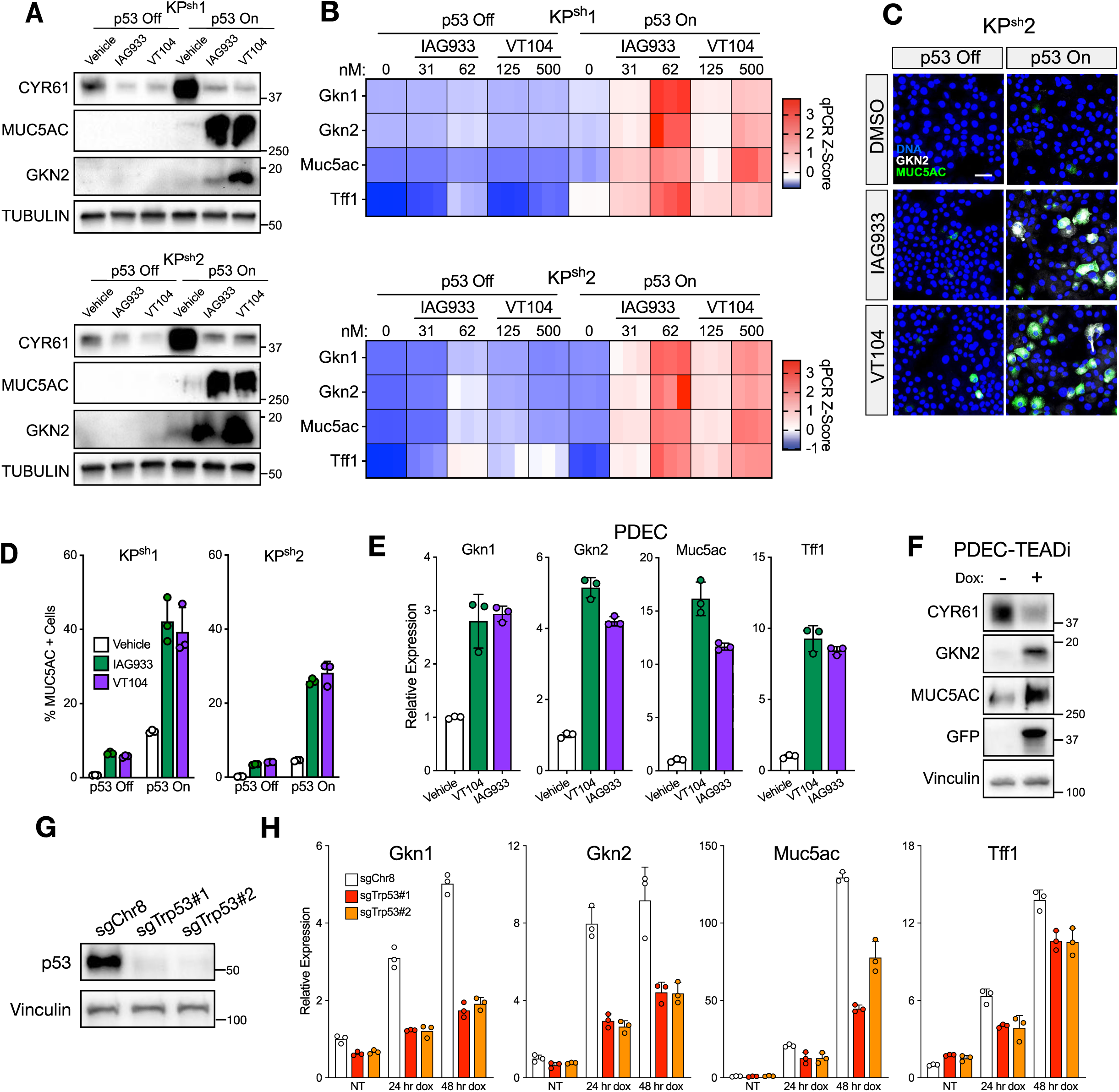
YAP/TAZ act through TEAD to repress gastric differentiation promoted by p53. **(A)** Western blots in cells treated with 48 hours of 125 nM IAG933, 500 nM VT104, or 1:1000 DMSO vehicle. **(B)** Heatmaps of RT-qPCRs of Z-scored expression with 48-hour treatments. **(C)** Representative 40x immunofluorescence images of GKN2 and MUC5AC in KP^sh^2 cells treated with 48 hours of 125 nM IAG933, 500 nM VT104, or DMSO vehicle. DNA was stained with Sytox Green. **(D)** Flow cytometry of MUC5AC showing percentage MUC5AC positive cells from 3 independent wells of KP^sh^ treated with 48 hours of 125 nM IAG933, 500 nM VT104, or 1:1000 DMSO vehicle. **(E)** RT-qPCRs from PDECs treated with 48 hours of 125 nM IAG933, 500 nM VT104, or 1:1000 DMSO vehicle. **(F-H)** PDEC cells expressing pInducer20-TEADi-GFP treated with 1 µg/mL dox for 48 hours. **(F)** Representative western blots of TEAD inhibition. **(G)** Western blots of bulk sgRNA-mediated knockout of Trp53. **(H)** RT-qPCRs from indicated treatment conditions. Scale bar in (C) = 50 µm.

GPL differentiation is a feature of benign, Kras^Mut^ metaplasia harboring wildtype p53 and is progressively lost following p53 inactivation during PDAC development^12,30^. Therefore, we next asked if acute loss of p53 would reduce the ability of TEAD inhibition to increase GPL marker expression in pre-malignant Kras^Mut^; p53 WT mouse pancreatic ductal epithelial cells (PDECs)^52,53^. Treatment of PDECs with IAG933 and VT104 (Figure 3E) or expression of a GFP-linked, dox inducible short peptide that prevents TEAD binding to YAP/TAZ in dominant negative fashion^54^ (Figure 3F, Supplemental Figure 2D,E) upregulated Gkn1, Gkn2, Muc5ac, and Tff1 expression, thereby demonstrating potent repression of GPL marker genes by TEAD activity in these cells. Consistent with a role for p53 activity in promoting gastric differentiation, CRISPR mediated knockout of p53 blunted the upregulation of Gkn1, Gkn2, Muc5ac, and Tff1 triggered by TEAD inhibition (Figure 2G,H). Therefore, YAP/TAZ acts through TEAD activity to suppress gastric differentiation driven by p53 activity in Kras^Mut^ pancreatic epithelial cells.

### Physiological and therapeutic adaptation connect YAP/TAZ activity to gastric fate in pancreatic cancer cells

YAP/TAZ levels are determined by a diverse set of cell intrinsic and extrinsic cues that are reflected in PDAC development and in cancer cell therapeutic response^43^. Given the key role that cell stress, cell-microenvironment, and cell-cell interactions play in dictating plasticity and transcriptional heterogeneity in PDAC development^55^, we tested if physiological cues that control YAP/TAZ levels also determine gastric differentiation outputs in the context of p53 activity.

Mechanical cues and cytoskeletal dynamics, including actin polymerization, can lead to YAP/TAZ activation^56,57^, leading us to test if changes in the actin cytoskeleton could play a role in YAP/TAZ-mediated control of gastric differentiation. Consistent with increased accumulation of actin stress fibers upon cell cycle arrest and entry into senescence^58^, p53 restoration resulted in prominent actin polymerization coincident with accumulation and nuclear localization of YAP/TAZ (Figure 4A). Remarkably, inhibiting actin polymerization via treatment with latrunculin-B both reduced YAP/TAZ nuclear accumulation and induction of Cyr61 in response to p53 restoration and resulted in upregulation of GPL markers Gkn1, Gkn2, Muc5ac, and Tff1 (Figure 4A,B, Supplementary figure 3A). Alternatively, impairing p53 induced senescence by silencing the p53 target gene p21/Cdkn1a in KP^sh^ cells reduced actin accumulation and total and nuclear TAZ following p53 restoration (Supplementary figure 3B-D) and similarly increased expression of GPL markers (Supplementary figure 3E). Thus, actin dynamics linked to cell cycle arrest and senescence connect YAP/TAZ activity to gastric gene expression in response to p53 activation.

**Figure 4.**
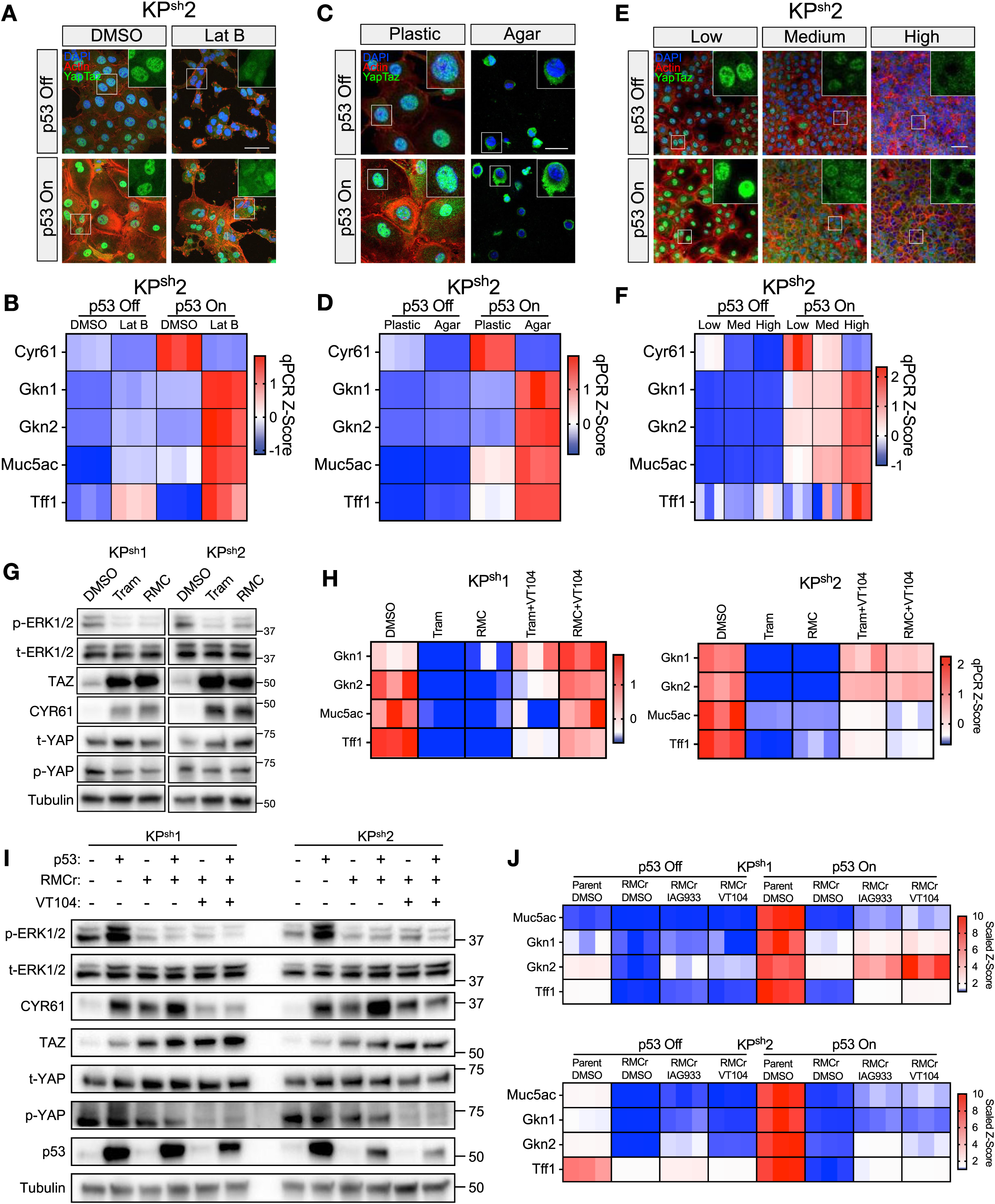
Physiological and therapeutic cues of YAP/TAZ activity dictates gastric differentiation. (A,B) KP^sh^2 cells treated with 12 hours of 5 µM Latrunculin B. **(A)** IF microscopy images stained with DAPI, Phalloidin (Actin), or YAP/TAZ antibody; scale bar 50 µm. **(B)** Heatmaps of RT-qPCR Z-scored expression. **(C,D)** KP^sh^2 cells seeded into 0.5% soft agar or on Collagen I coated plastic dishes and collected 24 hours after seeding. **(C)** IF microscopy images stained with DAPI, Phalloidin (Actin) or YAP/TAZ antibody; scale bar 25 µm. **(D)** Heatmaps of RT-qPCR Z-scored expression. **(E,F)** KP^sh^ cells were seeded at three different cell densities. **(E)** IF microscopy images of KP^sh^2 stained with DAPI, Phalloidin (Actin) or YAP/TAZ antibody; scale bar 50 µm. **(F)** Heatmaps of RT-qPCR Z-scored expression. **(G)** Representative western blots following treatment of 72 hours of 50 nM trametinib (Tram) or 50 nM RMC-7977 (RMC) in KP^sh^ grown on dox. **(H)** Heatmaps of RT-qPCR data from cells treated with 72 hours of 50 nM Tram or 50 nM of RMC followed by an additional 48-hour co-treatment with 500 nM VT104. **(I.J)** KP^sh^ parental lines or lines resistant to 100 µM RMC-7977 (RMCr). **(I)** Representative western blots with 48-hour 500 nM treatment of VT104. **(J)** Treatment with 48 hours of 125 nM IAG933, 500 nM VT104, or DMSO showing heatmap of RT-qPCR Z-scores scaled 1-10 of expression.

Cell growth in hyperconfluent cultures or soft matrices inhibits YAP/TAZ through cytoplasmic sequestration and has been shown to alter differentiation programs associated with YAP/TAZ function^56,59^. Given that microenvironmental stiffness can dictate YAP/TAZ accumulation in PDAC^60^, we next asked if changing local stiffness by altering growth strata or cellular packing can dictate YAP/TAZ mediated changes in GPL markers. Standard culture of KP^sh^ cells on Collagen I coated plastic results in largely nuclear YAP/TAZ localization, while seeding into a soft matrix of 0.5% Agar for 24 hours produced striking nuclear YAP/TAZ exclusion corresponding with reduced Cyr61 expression and Gkn1, Gkn2, Muc5ac, and Tff1 upregulation (Figure 4C,D, Supplementary figure 3F). Additionally, seeding of KP^sh^ cells at increasing cell densities resulted in a shift from high nuclear YAP/TAZ signal at low densities to near complete loss of YAP/TAZ nuclear localization in hyperconfluent cultures (Figure 4E). Cyr61 expression tracked inversely with cell density with upregulation of Gkn1, Gkn2, Muc5ac, and Tff1 in a density-dependent manner in cells following p53 restoration (Figure 4F, Supplementary figure 3G). Taken together, we find that gastric differentiation is intimately associated with YAP/TAZ levels in pancreatic cancer cells, that can be influenced by dynamic, microenvironmental associated factors influencing actin polymerization, extracellular stiffness, and spatial interactions.

Adaptive YAP/TAZ-TEAD activity is also emerging as a resistance mechanism to inhibition of the Kras/MAPK pathway^23,45,61^, a key therapeutic target in PDAC given the high frequency of KRAS mutations in the disease^62^. Furthermore, transcriptional heterogeneity along the classical-basal axis is implicated in response and resistance to emerging RAS inhibitors^63,64^. Therefore, we asked if changes in YAP/TAZ-TEAD activity in response to both short term Kras^Mut^ inhibition and Kras/MAPK inhibitor resistance connects with gastric differentiation. As previously demonstrated in human PDAC cells^43^, treating p53 silenced KP^sh^ cells with the pan-RAS inhibitor RMC-7779 (RMC) or the FDA approved MEK inhibitor Trametinib (Tram) for 72-hours resulted in robust TAZ stabilization, modestly reduced p-YAP, and elevated CYR61, confirming YAP/TAZ-TEAD activation (Figure 4G). This corresponded with potent inhibition of GPL genes Gkn1, Gkn2, Muc5ac, and Tff1, which was rescued via TEAD inhibition with VT104. (Figure 4H). Accordingly, RNA-seq analysis of KP^sh^ cells following 72 hours of Tram treatment revealed upregulation of a YAP/TAZ target gene signature and potent repression of the metaplastic gastric pit-like gene signature (Supplementary figure 4A).

Previous work has demonstrated that acquired resistance to RAS inhibition in PDAC leads to the loss of the classical subtype expression program^23,64^. We generated KP^sh^ cells resistant to RMC after long term treatment (RMCr) (Supplemental Figure 4B), which corresponded with elevated actin polymerization and increased levels of nuclear YAP/TAZ (Supplemental Figure 4C). Additionally, we observed high levels of CYR61 that were maintained in the setting of reduced phospho-ERK1/2^65^ (Figure 4I). Consistent with YAP/TAZ-TEAD mediated repression of GPL differentiation in RMCr cells, Gkn1, Gkn2, Muc5ac, and Tff1 were strongly downregulated and MUC5AC positivity was abrogated when compared to parental KP^sh^ lines and partially rescued by TEADi (Figure 4J, Supplemental Figure 4D). These data indicate that YAP/TAZ-TEAD activation in response to Kras/MAPK inhibition can repress gastric/classical differentiation and that this repression corresponds with the duration and degree of YAP/TAZ-TEAD activity.

### p53 and YAP/TAZ-TEAD activity interact to determine gastric heterogeneity *in vivo*

Restoring p53 in orthotopic murine PDAC promotes epithelial differentiation and features of pre-malignant differentiation *in vivo*^31^. Therefore, we asked if competition between p53 and YAP/TAZ-TEAD activity dictates gastric differentiation during tumor maintenance. First, we implanted KP^sh^-1 cells expressing shRNAs targeting YAP or TAZ orthotopically and randomized these mice into groups maintained on dox or normal chow (Supplemental Figure 5A). To confirm that shRNA-mediated knockdown of TAZ and YAP was maintained during tumor growth and p53 restoration, we sorted mKate2+ tumor cells following dox withdrawal and measured TAZ and YAP expression.

Expression analysis revealed that Wwtr1 (TAZ) knockdown was maintained in shTAZ tumors (Supplementary figure 5B), while YAP knockdown was not (Supplementary figure 5C). We hypothesize the differential maintenance of TAZ and YAP knockdowns likely owes to more modest YAP knockdown efficiency (Figure 1E) and increased essentiality for YAP over TAZ in the development and maintenance of PDAC^42,66^. While Taz silenced tumors did not display noticeable histological differences compared with controls (shRen) when p53 remained off, restoration of p53 in shTAZ tumors produced a dramatic increase in glandular structures containing cells with goblet-like morphology and large hollow apical structures consistent with mucin granules (Supplementary figure 5D). Immunofluorescence for GKN2 and MUC5AC confirmed that goblet morphology corresponded with GPL differentiation (Supplementary Figure 5D, E), thus demonstrating the ability of TAZ to repress gastric differentiation in the setting of p53 activity.

Next, we set out to test if the repression of gastric-like differentiation *in vivo* is mediated by TEAD transcriptional activity. First, we constitutively expressed the GFP linked, dominant negative TEADi peptide in KP^sh^ cells. Likely due to a role for TEAD activity in PDAC maintenance, we were only able to maintain modest TEADi-GFP expression in 1 of 2 KP^sh^ cell lines. However, upregulation of Gkn1, Gkn2, Muc5ac, and Tff1, largely in response to p53 restoration, was observed in this setting (Supplementary Figure 6A,B). We next generated an inducible TEAD inhibitor system that could be controlled independently of p53 expression. We leveraged the tamoxifen-inducible modified estrogen receptor (ERT2) that permits stabilization and nuclear translocation upon binding with 4-hydroxytamoxifen (4OHT)^67^. We placed ERT2 in frame with the TEAD binding domains (TBD) of VGLL4, YAP, and TAZ on the N-terminal or C-terminal end coupled with a FLAG tag for tracking expression, generating “TEADi-ERT2 (TE)” and “ERT2-TEADi (ET)” as well as a FLAG tagged ERT2 as a control “(E)” (Figure 5A).

**Figure 5.**
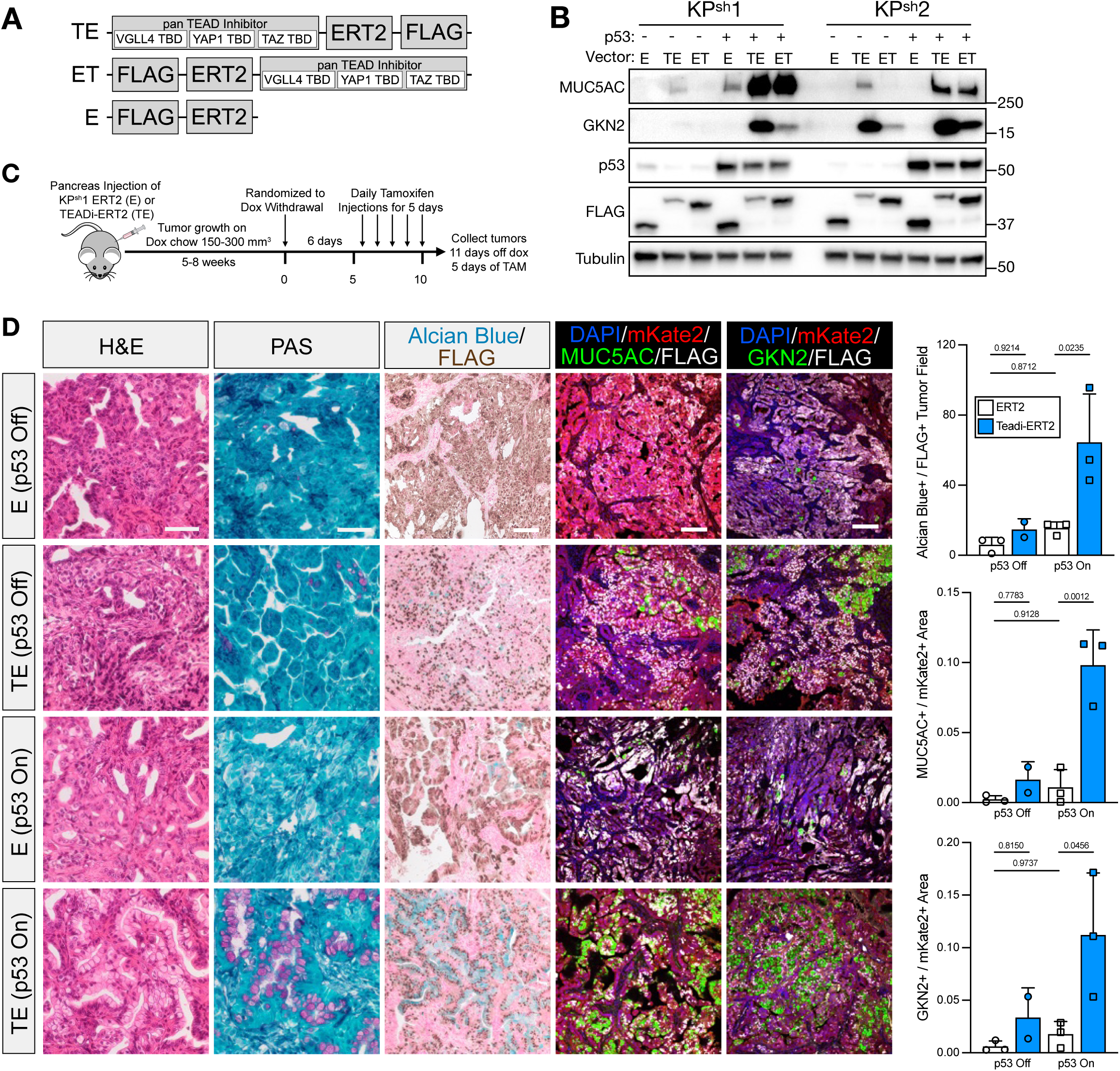
TEAD activity represses gastric differentiation driven by p53 *in vivo.* **(A)** Schematic demonstrating novel tamoxifen inducible dominant negative TEAD inhibitors. **(B)** Representative western blots of cells treated with 48 hours of 1 µM 4OHT. **(C)** Treatment regimen of orthotopic KP^sh^1 pancreas tumors for TEAD inhibitor induction. **(D)** Staining of orthotopic KP^sh^1 pancreas tumors on dox (p53 off) or 11 days off dox (p53 on) with tamoxifen regimen from (C). Column 1: Hematoxylin and Eosin (H&E) staining; Column 2: Periodic Acid Schiff (PAS) Staining; Column 3: Alcian blue Staining coupled with Immunohistochemistry using a FLAG antibody; Column 4 and 5: IF microscopy with antibodies against mKate2, FLAG, MUC5AC, and GKN2 with DAPI counterstain. Column 6: Quantification of area of Alcian blue positive cells per FLAG positive tumor area (top) or MUC5AC positive area (middle) and GKN2 positive area (bottom) per mKate2 positive area. Each dot represents quantification of one tumor from at least five randomly selected FLAG+ tumor fields. P-values calculated with one-way ANOVA with Tukey’s multiple comparisons test. Scale bars = 100 µm

4-OHT treatment caused robust stabilization of all constructs resulting in CYR61 downregulation in TE and ET expressing KP^sh^ cells (Supplementary Figure 6C). TE and ET localized to the nucleus while E was detected in the nucleus and cytoplasm (Supplementary figure 6D). Like treatment with pharmacological TEAD inhibitors or Yap/Taz knockdown, TEAD target genes Cyr61, Ctgf, and Thbs1 were downregulated in TE and ET expressing cells following tamoxifen treatment in p53 silenced and reactivated cells (Supplementary Figure 6E). To confirm TE and ET were interacting with TEAD, we performed immunoprecipitation (IP) with an anti-FLAG antibody revealing robust binding of TEAD1 to both TE and ET regardless of p53 status, while no TEAD1 was detected in E immunoprecipitant (Supplementary Figure 6F). 4-OHT treatment of TE and ET expressing cells following p53 restoration resulted in robust upregulation of GPL markers Gkn1, Gkn2, Muc5ac, and Tff1 (Supplementary figure 6G) and emergence of MUC5AC and GKN2 positive cells consistent with the effect of small molecule TEAD inhibitors and silencing of YAP/TAZ (Figure 5B, Supplementary figure 6H).

Using this system we next tested whether inducible TEAD inhibition could interact with p53 restoration to dictate gastric differentiation *in vivo*. KP^sh^ cells expressing E or TE were orthotopically injected into the pancreas of mice on dox chow. After tumor development, mice were randomized to stay on dox chow or undergo a 6 day dox withdrawal period, each followed by five daily consecutive doses of tamoxifen in all mice before harvest (Figure 5C). Histological analysis of TE tumors following p53 restoration revealed a dramatic increase of ductal structures with prominent goblet-like differentiation containing a hollow apical cytoplasm (Figure 5D). Periodic Acid Schiff (PAS) and Alcian blue staining confirmed extensive mucin granule formation containing acidic and neutral mucins, which was enriched in cells with detectable nuclear TE marked by FLAG following p53 activation (Figure 5D). Although Alcian blue, MUC5AC, and GKN2 were detectable in some FLAG+ mKate2+ cells in tumors where p53 remained silenced, TE activation in the setting of p53 restoration resulted in significantly greater expression of MUC5AC, and GKN2 per tumor area following p53 restoration. Thus, TEAD acts to suppress the gastric differentiation program promoted by p53 activation *in vivo*, suggesting that competition between p53 and YAP/TAZ-TEAD activity can dictate gastric pit-like plasticity during pancreatic cancer maintenance.

### p53 and TEAD activities predict GPL heterogeneity during PDAC development

Inactivation of p53 in Kras^Mut^ cells triggers the premalignant to malignant switch during which benign metaplastic cells adopt aggressive, invasive cell states^26^. Consistent with our observations, previous reports suggest that p53 wildtype metaplasia exhibits higher gastric gene expression than p53 inactivated, invasive disease^30^. To explore the relationship between p53 and YAP/TAZ-TEAD regulated gastric differentiation in PDAC progression, we leveraged the KPC^LOH^ model that permits tracking of cells that have undergone p53 loss of heterozygosity (LOH) based on a GFP reporter linked in cis with WT p53^26^ (Figure 6A). Here, expression of mutant Kras and deletion of one copy of p53 is marked by conditional expression of mKate2, allowing isolation of cells based on p53 status whereby pre-malignant double positive cells (DP, mKate2+, GFP+) harbor WT p53 while single positive cells (SP, mKate2+, GFP-) proceed to malignancy following sporadic loss of heterozygosity^26^.

**Figure 6.**
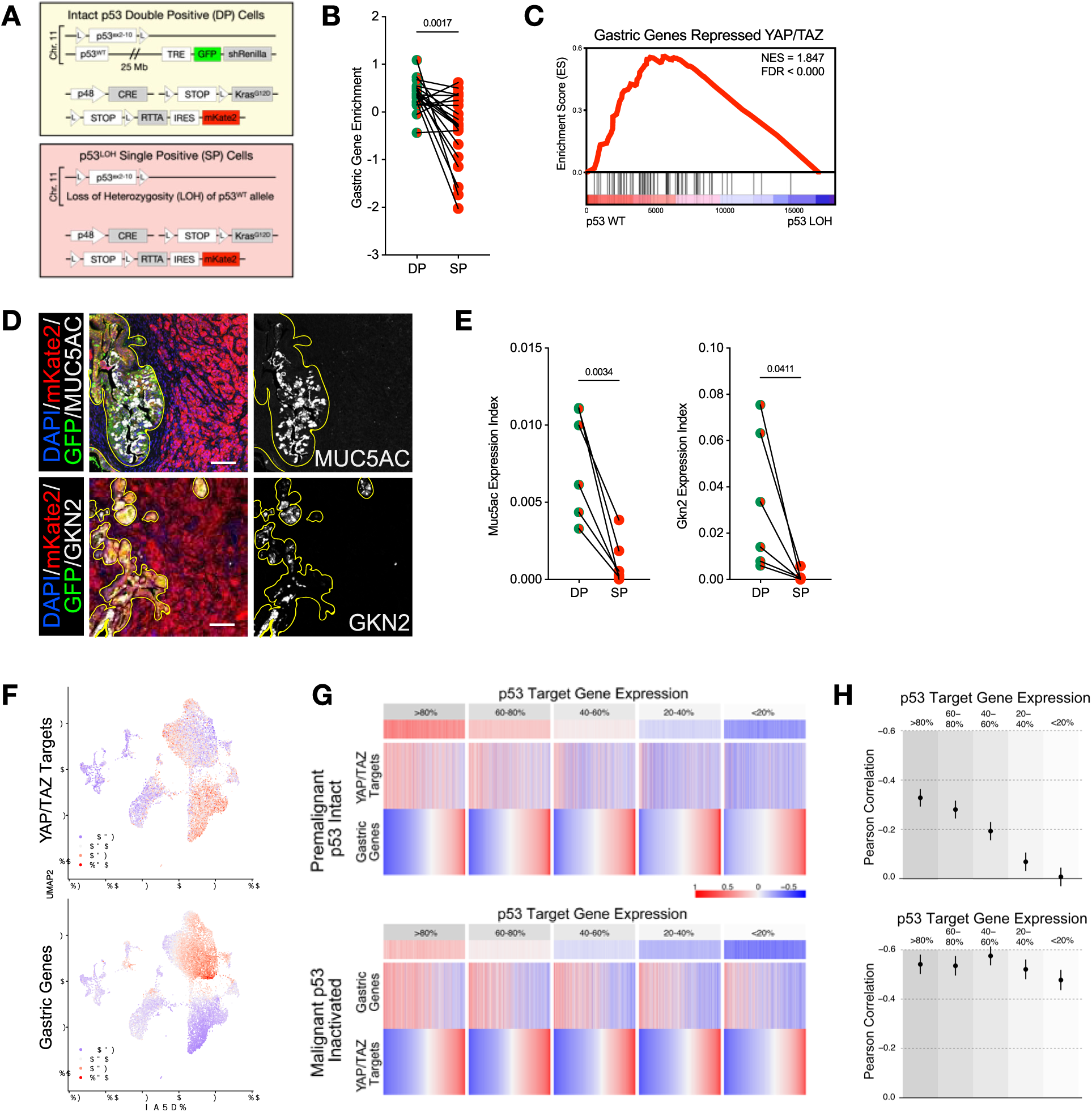
p53 and YAP/TAZ-TEAD activity dictate gastric heterogeneity during PDAC development. **(A)** KPC^LOH^ schematic **(B)** Enrichment of gastric genes from tumor matched DP and SP populations. P-value from paired Student’s T-test. **(C)** GSEA of from tumor matched DP and SP populations. **(D)** IF microscopy staining of SP:DP interface for MUC5AC (top) and GKN2 (bottom). **(E)** Quantification of IF microscopy images from (D) with each set of paired dots representing one SP:DP matched tumor from at least five fields. P-value from paired Students T-test. **(F)** UMAPs of scRNA-seq from K1-K4 Kras^mut^ metaplastic pancreas cells projecting YAP/TAZ target gene expression (top) or gastric genes (bottom). **(G-H)** scRNA-seq data from (top) p53 wild type Kras^mut^ cells (K1-K4) and (bottom) p53 inactivated Kras^mut^ pancreas cancer cells (K5-K6). **(G)** Heatmaps of gene expression program enrichment. **(H)** Metaplots depicting Pearson correlation between the expression scores for YAP/TAZ targets and gastric genes. Error bars showing 95% confidence interval.

We used FACS to isolate matched DP and SP cells from tumor bearing KPC^LOH^ mice and subjected them to RNA-seq. DP cells had significantly higher enrichment of gastric genes with 16/20 matched DP/SP pairs exhibiting reduced gastric gene enrichment upon p53^LOH^ (Figure 6B). GSEA on all tumors confirmed enrichment of the gastric genes in DP cells with nearly all YAP/TAZ controlled gastric genes more highly expressed in WT p53 populations (Figure 6C). Staining for MUC5AC and GKN2 confirmed abundant gastric marker expression in DP cells, which was largely absent in adjacent SP tumor areas (Figure 6D,E). These data highlight the role that p53 plays in enabling gastric differentiation during PDAC progression.

To explore the effect of p53 and YAP/TAZ-TEAD activity on gastric heterogeneity during PDAC development we leveraged previously published scRNA-seq profiling data reflecting p53 intact metaplasia and p53 inactivated mouse PDAC^27^. Projecting the expression score of a curated YAP/TAZ target gene signature^68^ and gastric genes repressed by YAP/TAZ (Figure 1) across Kras^mut^, p53 WT cells revealed a remarkable inverse correlation between populations with high gastric expression and high YAP/TAZ target expression (Figure 6F). Next, to investigate if p53 activity influences the YAP/TAZ-gastric fate anticorrelation, we added p53 inactivated PDAC cells to our analysis. Cells were ranked into five equal-sized bins based on enrichment of a p53 target gene signature, as an indicator of p53 activity, expression of YAP/TAZ targets, and markers of gastric pit-like identity (Figure 6G). In p53 WT cells, the Pearson anticorrelation between YAP/TAZ targets and gastric genes was highest in cells with the most p53 activity with a stepwise reduction as p53 activity fell (Figure 6H). However, although YAP/TAZ activity was anticorrelated with gastric gene expression in p53 inactivated PDAC, p53 target expression had little impact on the relationship. Indeed, fitting a linear model to predict gastric gene expression based on p53 and YAP/TAZ targets revealed a strong interaction in constraining gastric genes in p53 WT cells (interaction term= −0.59, p= 5.7e^-56^) while expression of p53 targets had little effect in p53 inactivated cells (interaction term= −0.14, p= 0.0087). Thus, while YAP/TAZ activity is associated with suppression of gastric fate regardless of p53 status, gastric heterogeneity in Kras^Mut^ pre-malignant metaplasia is strongly determined by the interplay between p53 and YAP/TAZ-TEAD activity.

### YAP/TAZ-TEAD activity constrains gastric and classical subtype differentiation in PDAC

Given the ability of YAP/TAZ-TEAD activity to suppress gastric and classical subtype gene expression in mouse pancreatic cancer cells, we asked if this regulatory interaction extended to human PDAC. Consistent with mouse PDAC, bulk expression analysis from four cohorts of PDAC patients (n=626) revealed an anti-correlation between YAP/TAZ-TEAD targets and both gastric pit-like differentiation (r=-0.50, p=1.1e-41) as well as classical subtype gene expression (r=-0.397, p=4.2e-25) (Figure 7A). Like the spectrum of metaplastic heterogeneity observed in mouse models, single cell expression analysis of 200 patient samples revealed a marked inverse correlation between GPL differentiation and key markers of the classical transcriptional subtype with YAP/TAZ-TEAD activity signatures (Figure 7B). To confirm that TEAD activity directly represses the gastric fate in human PDAC, we treated a panel of 11 human PDAC cell lines with the TEAD inhibitor IAG933. TEAD inhibition downregulated YAP/TAZ-TEAD targets (CTGF, CYR61) and resulted in strong enrichment of gastric genes (MUC5AC, TFF1) and markers of the classical subtype (LGALS4, LYZ, etc) (Figure 7C). TEAD inhibition increased the frequency of MUC5AC positive cells via both IF staining and flow cytometry (Figure 7D,E, Supplementary Figure 7A). Furthermore, activation of a dox-inducible dominant negative TEAD inhibitor upregulated gastric/classical genes and increased MUC5AC positive HPAC cells in a dose-dependent manner (Supplementary Figure 7B-D). These data validate a conserved role for YAP/TAZ-TEAD in repression of gastric differentiation overlapping with the classical subtype in human PDAC.

**Figure 7.**
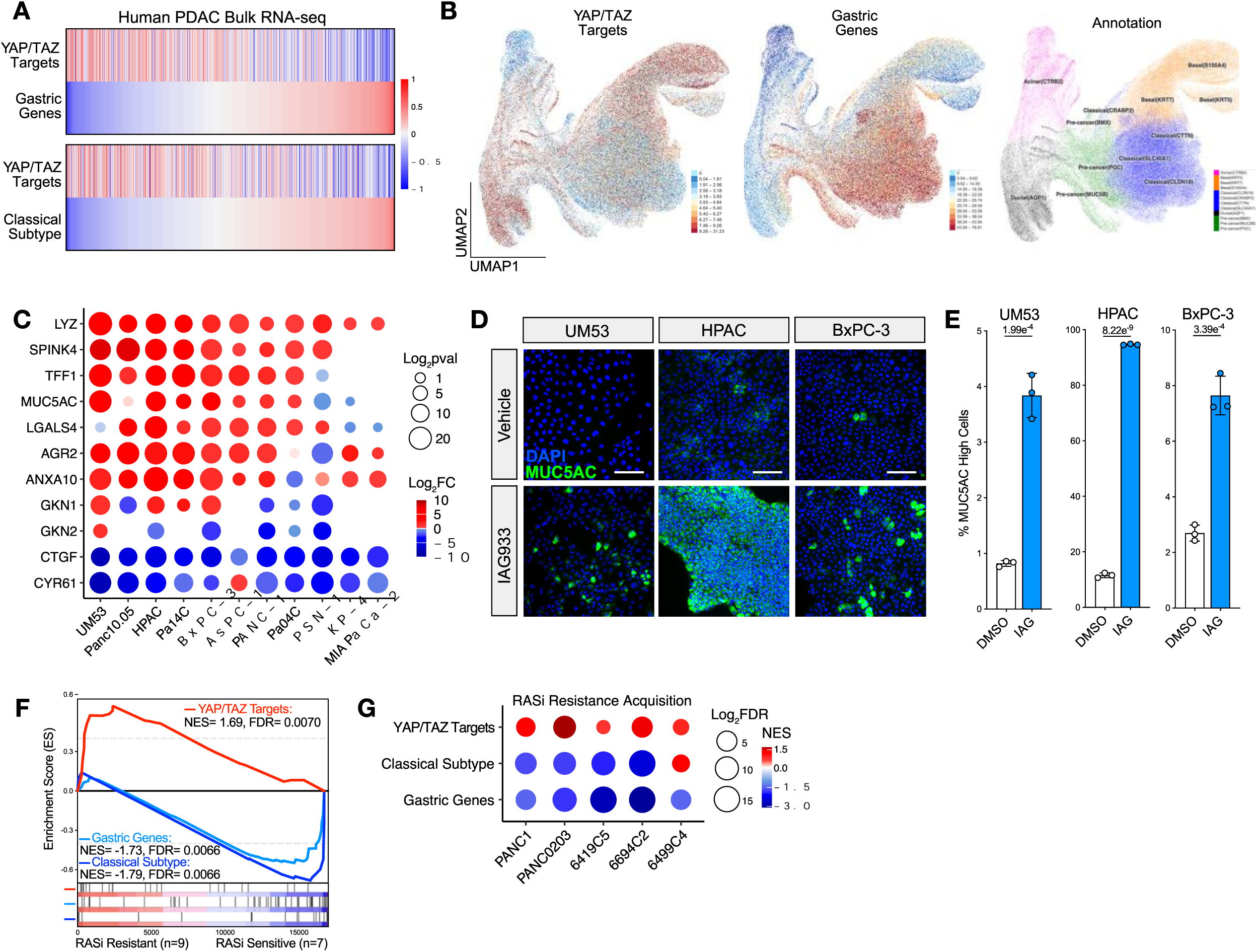
YAP/TAZ-TEAD represses gastric and classical subtype differentiation in PDAC. **(A)** Bulk RNA-seq of n=626 human PDACs showing scaled gene signature enrichment scores for each tumor. **(B)** scRNA-seq of n=200 human PDACs projecting enrichment of (Left) YAP/TAZ targets and (Middle) gastric genes with (Right) associated subtype classification annotations. **(C)** qRT-qPCR, **(D)** representative IF microscopy images, and **(E)** percent MUC5AC positivity in a panel of human PDAC cell lines treated with 48 hours of 1 µM IAG933. **(F)** GSEA from RNA-seq of cells stratified by MRTX1133 sensitivity^64^. **(G)** GSEA on mouse (n=3) and human (n=2) PDAC lines comparing parental cells and cells exhibiting acquired resistant to MRTX1133 by dose escalation^64^. Scale bar in (D) = 50 µm

Both innate/primary and acquired resistance to RAS inhibition has been linked with elevated YAP/TAZ activity in human PDAC cell lines^23,45,61^. Reanalysis of RNA-seq data from human PDAC lines stratified into sensitive (n=7) or innately resistant (n=9) to the Kras^G12D^ inhibitor MRTX1133 (MRTX) revealed enrichment of YAP/TAZ targets in MRTX resistant cells associated with a deficiency in gastric fate and classical subtype genes^64^ (Figure 7F). Furthermore, acquired MRTX resistance by dose escalation in 2 human and 3 mouse PDAC cell lines facilitated YAP/TAZ target upregulation and loss of the classical subtype and gastric gene expression (Figure 7G). Therefore, adaptive YAP/TAZ-TEAD activation in response to RAS inhibitors can dynamically regulate gastric and classical subtype differentiation in PDAC.

## DISCUSSION

Despite a limited set of recurrent cancer driving mutations, PDAC displays pervasive molecular heterogeneity that manifests as a spectrum of plastic cellular identities during both pre-malignant initiation and cancer progression. It is well established that the 2 most common PDAC driver events, mutations in Kras and inactivation of p53, act to remodel pancreatic plasticity during PDAC development. Mutant Kras triggers metaplastic reprogramming of acinar cells into a molecularly heterogenous, duct like lineage characterized by persistent activity of pathways that play key, temporally regulated roles in development and regeneration. Inactivation of p53 in these pre-malignant cells triggers progression to malignancy and the erosion of well differentiated metaplastic cell fates. However, how genetic drivers of PDAC interact with aberrantly activated signaling pathways to determine plastic, cellular identities remain poorly understood. Here, using a mouse model which permits restoration of wildtype p53 in advanced pancreatic cancer cells, we demonstrate that the activity of p53 and YAP/TAZ-TEAD transcriptional complexes compete to determine the expression of a gastric pit-like (GPL) differentiation program observed during pre-malignant specification that is also characteristic of the classical PDAC transcriptional subtype. We find that p53 enables expression of GPL differentiation while YAP/TAZ-TEAD broadly represses it. GPL differentiation is exquisitely linked to YAP/TAZ-TEAD activity as both genetic and physiological mediators of YAP/TAZ levels, including cellular adaptation to potent inhibition of RAS/MAPK signaling, can determine GPL/classical marker gene expression. This suggests that both microenvironmental cues and states of cellular stress that regulate Hippo signaling likely influence metaplastic heterogeneity. Indeed, analysis of mouse and human single cell expression data sets demonstrate a consistent, inverse correlation between YAP/TAZ-TEAD activity signatures and GPL/classical differentiation in mouse and human PDAC, which we recapitulate by independently regulating p53 and TEAD activity in pancreatic tumors *in vivo*. Inhibiting TEAD in human PDAC cells derepresses GPL and classical differentiation marker genes while accumulation of YAP/TAZ in response to acute and long-term inhibition of RAS/MAPK signaling in mouse cancer cells actively suppresses GPL/classical differentiation. Taken together, our work highlights the ability of two key determinants of PDAC progression, p53 and YAP/TAZ-TEAD, to determine GPL and classical heterogeneity throughout PDAC development and in response to targeted therapies.

Several reports have identified gastric pit-like fate as a prominent cellular identity in heterogeneous pre-malignant metaplasia via scRNA-seq and histological analysis^12,25,27,30^. Acquisition of the gastric pit-like program is specifically associated with Kras^Mut^ with markers of GPL largely found in PanINs^11,13,30^. As confirmed by our innovative lineage tracing approach, upon p53 inactivation the gastric fate becomes less accessible, especially in aggressive, poorly differentiated disease^26,30^. Restoring p53 activity in KP^sh^ PDAC cells is sufficient to upregulate GPL marker expression, while deleting p53 in pre-malignant Kras^mut^ duct cells blunts GPL marker genes, consistent with a role for p53 in promoting epithelial differentiation programs and the colocalization of gastric and senescent cells within the same PanINs^12^. Interestingly, p53 restoration also increases TAZ levels in cells, which acts to limit GPL marker gene expression suggesting a framework whereby p53 acts at multiple levels of molecular control to determine neoplastic fate along the ductal-gastric axis in PDAC development.

Our work identifies YAP/TAZ-TEAD activity as a negative regulator of gastric differentiation in PDAC development. In normal tissue, YAP/TAZ similarly restrict goblet differentiation and mucin production genes in the airway epithelium via SPDEF repression^69^. Interestingly, SPDEF also drives a mucinous production program in pancreatic metaplasia and Nkx6-2 promotes gastric differentiation in intraductal papillary mucinous neoplasms^30,70^. Similar gastric programs are observed in Kras^Mut^ lung adenocarcinoma driven by FoxA1/2-TET DNA demethylation^71,72^. These data support a model of complex interplay of lineage defining transcription factors co-opted across organs to influence gastric-like heterogeneity in both endoderm development and tumorigenesis.

The classical PDAC subtype is enriched for gastric-like differentiation genes and exhibits differential therapeutic responses including greater sensitivity to traditional chemotherapy but resistance to novel RAS inhibitors. Acute RAS inhibition in tumors potentiates classical subtype gene expression consistent with reduced YAP/TAZ activity following short term RASi in vivo^23,61^. However, tumor relapse from long term RASi results in YAP/TAZ activation as a resistance mechanism corresponding with near complete loss of the classical program^23,45,61,64^. Here, we demonstrate that adaptive accumulation of YAP/TAZ dictates GPL and classical differentiation after acute and long term RAS/MAPK inhibition. We show that rapid accumulation of YAP/TAZ following RAS/MAPK inhibition suppresses GPL and classical differentiation markers. Furthermore, we validate that acquired resistance to the pan-RAS inhibitor RMC-7779 results in sustained YAP/TAZ accumulation and dramatic loss of GPL/classical differentiation. Critically, suppression of GPL and classical differentiation can be partially rescued by inhibition of TEAD at both timepoints. Considering that classical/basal heterogeneity underlies differential therapeutic susceptibilities, our work has implications for combining pharmacological TEAD inhibitors with other therapeutics to maximize the therapeutic benefit of skewing metaplastic heterogeneity.

## Methods

### Cell Culture

Primary KP^sh^ cells were obtained as previously described^31^ and plated onto dishes coated with Type I Bovine Collagen Solution (PureCol, Advanced Biomatrix, 0.1 mg/mL) in DMEM containing 10% tetracycline-free fetal bovine serum (FBS, Gibco) and 1x penicillin-streptomycin (P/S, Sigma). 1 μg/mL dox was added into medium during passaging (p53 off) or removed for 6 days to restore p53 expression (p53 on) unless otherwise stated. To generate RMC-7779 resistance, (RMCr) KP^sh^ cells were generated by culturing in the presence of 100 nM RMC-7779 and 1 μg/mL dox for at least 3 weeks until surviving cells resumed proliferating. RMCr KP^sh^ were maintained on dox and 100 nM RMC while parental KP^sh^ were passage matched with DMSO. Primary Pancreatic Ductal Epithelial Cells (PDECs) were derived as previously described (Pylayeva-Gupta et al. 2013). PDEC culture plates were first incubated with a low volume of 1:100 dilution of matrigel in DMEM for 1 hour at 37 degrees C before addition of pancreatic medium (PM) containing 10 ng/mL mouse EGF (Gibco), 400 ng/mL Dexamethasone (Sigma), 5 nM 3,3’,5-Triiodo-L-thyronine (Sigma), 25 μg/mL Bovine Pituitary Extract (Sigma), 0.5x ITS+1 (Sigma), 100 ng/mL Cholera toxin (Sigma), 100 μg/mL Soybean Trypsin Inhibitor (VWR), 1x P/S, and 10% heat inactivated FBS (Gibco) in DMEM/F-12 50:50 1x (VWR). Human PDAC lines Pa04c, KP-4, PANC-1, Mia PaCa-2, and Pa14C were grown in DMEM with 10% FBS and 1x P/S and PSN-1, AsPc-1, BxPC-3, Panc10.05, HPAC, and UM53 were grown in RPMI with 10% FBS and 1x P/S. Cell lines were periodically confirmed mycoplasma free via the Venor™ GeM Mycoplasma Detection Kit (Sigma).

### Generating Tamoxifen inducible ERT2-TEADi

Sequences for ERT2 (E), TEADi-ERT2 (TE), and ERT2-TEADi (ET) (Table 2) were purchased as G blocks from Integrated DNA technologies. G blocks were digested with BamHI and EcoRI to create overhangs for ligation into pBABE-Puro (Addgene #21836). Retrovirus was prepared as above and stably transduced into KP^sh^ cells.

### Soft Agar Experiment

KP^sh^ cells were kept on dox or removed from dox for five days before trypsinization and seeding. For making the soft agar substrate, agar powder was diluted to 1% in DMEM+10% FBS, boiled in the microwave to dissolve, and a thin layer was added to culture dishes to solidify. Cells were then resuspended in 0.5% dissolved agar layered on top of the solidified 1% agar matrix. For RNA, the 0.5% agar matrix was scraped away from a 6-well plate and mixed with BME containing RLT buffer for lysis and compared to cells seeded onto a collagen-coated 6-well. For imaging, cells in an agar plug from a 12-well plate were compared with cells seeded in an Ibidi-Treated µ-well 4-chamber slide. Cells contained within the agar were fixed with 4% paraformaldehyde with 0.1% glutaraldehyde in PBS, washed with PBS, then incubated in 15% sucrose overnight at 4 degrees. The agar was embedded in OCT, frozen on dry ice, and cryosectioned onto charged slides. After three 10-minute incubations in 20 mM sodium borohydride in PBS, IF staining was performed as described above.

### Autochthonous KPC^LOH^ Tumor Model

*p48-Cre*, *LSL-Kras^G12D^*, *p53^flox^*, *CHC*, and *CAGs-LSL-RIK* strains were maintained on mixed Bl6/129J as described previously^26^ and grown on dox chow (625 mg/kg, Harlan Laboratories). Upon cancer formation detected by small animal ultrasound, pancreata were harvested for either formalin fixation for staining or dissociated for flow cytometry to isolate SP and DP populations. Staining was performed for six mice harboring SP cancers with matched adjacent DP premalignant metaplastic tissue.

### Orthotopic Pancreas Tumor Model

250,000 KP^sh^1 cells harboring shREN, shYAP#1, shTAZ#2, ERT2 control, or TEADi-ERT2 suspended in 15 μL of a 1:1 suspension of Matrigel:DMEM were injected into the pancreas of athymic nude mice placed on dox chow for 3 days prior. Small animal ultrasound was performed to track tumor growth until tumors were approximately 500 mm^3^ for shRNA tumors or 150-300 mm^3^ for the tamoxifen inducible TEAD inhibitor tumors. Animals were randomized to stay on dox or be removed from dox to restore p53 and tumors were harvested after 15 days or at humane endpoint (shRNA experiment) and 11 days (TEADi experiment). 6 days following randomization, all mice in the TEAD inhibitor cohort received daily intraperitoneal injections of 100 μL of 10 mg/mL tamoxifen dissolved in corn oil for ERT2 activation.

### RNA-seq of KP^sh^ cells

Two RNA-seq experiments in KP^sh^ were performed: KP^sh^1 cells on dox or 6 days off dox expressing constitutive shRNAs targeting Renilla, Yap1, or Wwtr1 and parental KP^sh^1/2 cells treated with 72 hours of 25 nM trametinib, vehicle on dox, or vehicle 6 days off dox. Triplicate wells of RNA were collected by scraping into RLT lysis buffer with BME, homogenized with a QIAshredder (Qiagen), and extraction with RNeasy mini kit (QIAgen). Novogene performed standard PolyA enrichment, cDNA preparation, and sequencing on a NovaSeq PE150. Raw data were processed with Partek Flow analysis software version 10.0.23.0720. Bases were trimmed for quality score > 20 and STAR (2.7.8a) aligned to the mm10 mouse genome. Gene counts were generated with HTseq followed by differential analysis by DESeq2 of all genes exhibiting the lowest maximum coverage of 2.0 counts. Gene Set Enrichment Analysis (GSEA version 4.3.2) was performed on median of ratios normalized counts using default parameters with 1000 gene set permutations.

### Mouse scRNA-seq data analysis

We analyzed scRNA-seq data from pancreatic metaplastic epithelial cells from stages K1-K6 from Burdziak et al. 2023 (accession number GSE207943)^27^. There are 13,650 pre-malignant cells from K1-K4, which are Kras-mutant and p53-wild type, and 8,855 malignant cells from K5-K6, which are Kras-mutant and p53-LOH. The UMI counts were normalized for library size and log-transformed using the standard Seurat pipeline^73^. We used the *AddModuleScore* function from the R package Seurat to compute expression scores of the gastric, YAP/TAZ and p53 gene signatures for each cell, and scaled the expression scores for each gene signature such that the scaled expression scores range from –1 to 1 across all cells. For each cell population (p53-wild type and p53-LOH), we divided all cells into 5 equal-sized groups based on the expression level of p53 target genes and plotted the heatmap showing the expression score of gastric, YAP/TAZ and p53 gene signatures for each group of cells, with cells ordered by the expression of gastric program within each group. To investigate whether and how the anti-correlation between gastric and YAP/TAZ expression scores depends on p53 activity, we generated a metaplot, separately for p53-wild type and p53-LOH populations, which displayed the Pearson correlation between gastric and YAP/TAZ expression scores and accompanying 95% confidence interval within each group of cells.

## Supplementary Methods

### Retrovirus and Lentivirus Production and Delivery of shRNA, sgRNA, and cDNAs

Virus was generated in HEK293Ts by transfecting cDNA, sgRNA, or shRNA cargo and packaging plasmids for virus using Lipofectamine 3000 (Thermo Scientific) according to manufacturer’s protocol. pBS-CMV-GagPol (Addgene #35614) and pCMV-VSVG (Addgene #8454) were used to make retrovirus while psPAX2 (Addgene #12260) and pMD2.G (Addgene #12259) were used to make lentivirus. Medium was changed 24 hours after transfection before beginning virus collection, filtering through 0.45 μm PES membrane, and infection for 48 hours in the presence of 16 μg/mL polybrene. shRNAs targeting Renilla luciferase (control), Cdkn1a (p21), Yap1, Wwtr1, and Lats1 were cloned into the retroviral LMNe-BFP vector. LMNe-BFP infected cells were selected using 1 mg/mL geneticin/G418. Knockout of Trp53, Yap1, Wwtr1, and a control intergenic region of Chromosome 8 (Chr8) was performed by stably introducing lentiCas9-blast followed by lentiviral delivery of sgRNAs cloned into pUSEPB backbone^31^. LentiCas9-blast selection and pUSEPB selection was performed with 10 μg/mL blasticidin for 7 days and 10 μg/mL puromycin for 7 days respectively. pInducer-20 EGFP-TEADi (Addgene, #140145) was transduced into PDEC and HPAC cells followed by five days of selection using 1 mg/mL geneticin/G418. cDNAs for wildtype and phospho-null YAP and TAZ and EGFP-TEADi were cloned out of pInducer-20 and into the retroviral backbone pBABE-puro for transduction into the KP^sh^ cells followed by selection with 10 μg/mL puromycin for 7 days^43^.

### Western Blotting

RIPA buffer (Thermo Scientific) supplemented with protease inhibitor cocktail (Roche) and phosphatase inhibitor (PhosSTOP, Roche) were used to lyse the cells. Protein concentrations were normalized via BCA, Laemmli buffer containing beta mercaptoethanol (BME) was added, and equivalent protein amounts were loaded onto SDS-PAGE gels. Proteins were transferred to 0.45 μm PVDF membranes (Immobilon P), blocked with 5% milk or BSA for total or phospho antibodies respectively for at least one hour at room temperature, then incubated overnight in primary antibodies diluted in blocking buffer at 4 decrees C. HRP conjugated secondary antibodies were added 1:10,000 in blocking buffer for one hour at room temperature. Clarity ECL (BioRad) was added to membranes for imaging on a chemidoc (BioRad).

### Real Time Quantitative Polymerase Chain Reaction (RT-qPCR)

RNA was extracted using RNeasy mini kit (Qiagen) according to manufacturer’s protocol. Briefly, cells were scraped into RLT lysis buffer containing BME, homogenized with QIAshredders (Qiagen), and processed through the RNEasy columns to isolate RNA. 1 μg of cDNA was synthesized using iScript reverse transcriptase (BioRad). RT-qPCR using PowerUp SYBR green (Thermo Fisher) was performed with a CFX Opus Real-Time PCR System (Bio-Rad). RT-qPCR primers were designed with MGH Harvard’s PrimerBank.

### Flow Cytometry Staining

Triplicate wells of KP^sh^ or human PDAC cells were seeded into 6-well plates. Cells were detached with TrypLE (Gibco) and washed with PBS before fixation and permeabilization with the Foxp3 Transcription Factor Fixation/Permeabilization kit (Invitrogen) on ice for one hour. Permeabilization (perm) buffer was used to wash cells twice before resuspending in perm buffer containing 2% FBS and 2% FC Block (Life Technologies, 14-9161-73). MUC5AC (45M1) (MA5-12178, Thermo Scientific) or Mouse mAb IgG Isotype control (Cell Signaling Technologies (E7Q5L), 53484S) primary antibodies were added directly to this solution at 1:100 overnight at 4 degrees. After washing in perm buffer, goat anti-Mouse IgG1 Alexa Fluor 647 Secondary Antibody (Invitrogen, A-21240) was added at 1:100. Cells were washed and subjected to flow cytometry using an Attune NxT flow cytometer. For gating, acellular debris were excluded using FSC-A vs SSC-A and doublet cells were removed with SSC-A vs SSC-H. Single cells were then gated as positive or negative for MUC5AC based on a conservative gate using the negative control.

### Immunofluorescence Microscopy

Cells were seeded into ibi-Treated 4 well µ-slides (Ibidi) coated with a PureCol (KP^sh^), Matrigel (PDEC) or uncoated (human PDAC lines). A 15-minute incubation at 37 degrees C in 4% paraformaldehyde was used for fixation and membranes were permeabilized with 0.3% Triton-X100. Cells were blocked using 5% BSA in PBST. Primary antibodies incubation was performed at 4 degrees C overnight in 3% BSA in PBST. Primary dilutions were as follows: 1:200 YAP/TAZ (D24E4) (CST, 8418), 1:250 TAZ (E9J5A) (CST, 72804), 1:350 YAP (D8H1X) (CST, 14074), 1:100 GKN2 (Invitrogen, PA5-119135), and 1:200 MUC5AC (45M1) (Thermo Scientific, MA5-12178). Secondary antibodies were incubated for 1 hour at room temperature protected from light in 5% BSA in PBST at 1:500 dilution. Following a quick wash, a 10-minute incubation was performed to stain actin with 1:2500 Phalloidin-iFluor 594 Reagent (Abcam, ab176757) and DNA with 1:30,000 SYTOX green in TBS (Thermo Fisher, S7020) or 1 μg/mL DAPI in PBS (Sigma, D9542). Imaging was performed with an Olympus IX81 or FV1000 Confocal Microscope (Olympus). Fiji (ImageJ) was used for image quantification.

### Growth and Viability Curves

KP^sh^1/2 expressing hairpins targeting Renilla luciferase or Cdkn1a were either kept on dox or removed from dox for 2 days for washout of the Trp53-targetting shRNA. 5,000 on dox and 15,000 off dox cells were then seeded into 12-well plates. Every 24 hours, cells were trypsinized and a Guava easyCyte was used to count triplicate wells. Fold change cell number is normalized to the number of cells counted 24 hours after seeding. For cell viability curves, 500 parental and 1000 RMCr KP^sh^1/2 were seeded into 96-well plates for 24 hours before the addition of RMC for a 96-hour incubation in 75 μL of drug containing medium. 60 μL of cell titer glo was added and luminescence was measured on a GloMax plate reader (Promega).

### Tumor Cell Fluorescence Associated Cell Sorting (FACS)

For flow, tumors were chopped finely, rinsed in PBS, and incubated on a rotator at 37 C with 1 mg/mL Collagenase V (Sigma, C9263) and 1 mg/mL Dispase II (Sigma, D4693) dissolved in PBS containing 0.1 mg/mL DNase I (Sigma, DN25). Cells were washed with FACS wash buffer (10 mM EGTA, 2% FBS, 0.1 mg/mL DNAse I in PBS), resuspended in 0.05% Trypsin-EDTA (Gibco) for 5 minutes at 37 C, washed, filtered through a 0.40 μm membrane, and resuspended in FACS collection buffer (10 mM EGTA, 2% FBS, 0.1 mg/mL DNAse I in DMEM), with 1 μg/mL DAPI for dead cell exclusion. For the shRNA orthotopic tumor cohort, acellular debris, doublet cells, mKate2 negative cells, and DAPI positive cells were excluded thereby sorting live, mKate2 positive singlet tumor cells directly in RLT buffer containing BME. RNA lysates were then processed as above for qPCR analysis of Yap1 and Wwtr1 knockdown. For the KPC^LOH^ cohort, GFP-/mKate2+ p53 LOH (SP) cancer cells and GFP+/mKate2+ p53 intact DP premalignant cells from the same mice were sorted using a FACS Aria III (UNC flow cytometry core facility) using the previously described gating strategy^26^. Sorted cells were centrifuged, resuspended in RLT with BME, homogenized with QIAshredders, and stored at −80C until RNA isolation using All prep RNA/DNA Micro Kits (QIAgen, 80284). Library preparation and RNA-seq was performed with Novogene’s HiSeq PE150 platform.

### Hematoxylin and Eosin (H&E) Staining

Tissues were deparaffinized and hydrated in xylenes and progressive ethanol dilutions ending with water. Tissues were incubated in Eosin Y (Sigma-Aldrich, Cat#318906), rinsed with water, and immersed in Mayer’s hematoxyin (Fisher Scientific, Cat#VMH032). A progressive ethanol series followed by xylenes were used for dehydration and clearing before mounting with VectaMount. 40x images were captured using an Olympus IX70 inverted microscope.

### Periodic Acid Schiff (PAS) Staining

Periodic Acid Schiff (PAS) Stain Pack (Tyr Scientific LLC, Cat#B083XQZXN2) was used for mucin visualization in tissues according to manufacturer’s recommended protocol. Briefly, tissue was deparaffinized and hydrated in xylenes and progressive ethanol dilution to water. Tissues were subjected to the PAS staining series via 7 minutes in PAS, 20 minutes in Schiff’s solution, 3 minutes in Mayer’s hematoxylin, 10 seconds in Bluing reagent, 2 minutes in light green solution, and a few seconds in ethanol with water washes between each staining step. Dehydration with graded ethanol and xylenes was followed by mounting in VectaMount (Vector). 40x Images were captured using an Olympus IX70 inverted microscope.

### Immunohistochemistry and Alcian Blue Staining

Tissues were deparaffinized in xylenes followed by progressive ethanol dilutions and washing in water. Tissue was pressure cooked for 15 minutes on high in a citrate-based antigen unmasking solution (Vector) followed quenching of endogenous peroxidases via incubation in 1% Hydrogen peroxide diluted in PBS at room temperature. 5% BSA in PBS was added to block for one hour followed by overnight primary antibody incubation with rabbit anti-FLAG clone D6W5B (Cell Signaling Technologies, Cat#14793S) at 1:200 dilution in blocking buffer. After washing in PBS, ImmPRESS® HRP Anti-Rabbit IgG (Peroxidase) Polymer (Vector) was added for one hour and ImmPACT™ DAB Peroxidase (HRP) Substrate (Vector) was used for FLAG visualization. Excess substrate was rinsed away with water and tissues were acidified in 3% acetic acid. Alcian Blue (pH 2.5) (Vector, Cat#H-3501) was added for 30 minutes at room temperature, washed briefly in 3% acetic acid, and rinsed with water. Nuclear Fast Red (Vector) was added for 5 minutes followed by a thorough water rinse. Tissue was then dehydrated through graded ethanol dilutions, cleared with xylenes, and mounted with VectaMount. 20x Images were captured using an Olympus IX70 inverted microscope for visualization.

### Tumor Immunofluorescence

Tissue was deparaffinized with xylenes and hydrated with progressive ethanol series of 100%, 95%, and 70% followed by water. Antigens were unmasked via pressure cooking for 15 minutes on high in a citrate-based antigen unmasking solution (Vector) and washed in PBS. Tissue was blocked with 5% BSA in PBS for at least one hour followed by overnight primary antibody incubation in blocking buffer with the following antibodies: 1:200 rabbit anti-FLAG (D6W5B) (Cell Signaling Technologies, Cat#14793S), 1:200 mouse anti-FLAG (M2) (Sigma-Aldrich, Cat#F3165), 1:200 rabbit anti-Gastrokine 2 (Invitrogen, PA5-119135), 1:300 mouse anti-Mucin 5Ac (45M1) (Thermo Scientific, Cat#MA5-12178), 1:500 anti-GFP (Abcam, 13970), and 1:100 of an in-house generated guinea pig anti-mKate2 antibody. Following PBS washing, secondary antibodies were added for at least one hour using a 1:500 dilution: Goat anti-Mouse IgG (H+L) Alexa Fluor™ 488 (Thermo, Cat#A-11001), Goat anti-Guinea Pig IgG (H+L) Alexa Fluor™ 568, (Thermo, Cat#A11075), and Goat anti-Rabbit IgG (H+L) Cross-Adsorbed Secondary Antibody, Alexa Fluor™ 647 (Thermo, Cat#A-21244). Following a quick wash, DAPI was added at 1 μg/mL for 15 minutes and Prolong Gold antifade reagent was added for mounting the coverslip. 20x images were captured with a FV1000 Olympus confocal microscope.

### Tumor Staining Quantification

For alcian blue quantification, at least ten randomly selected 40x fields containing FLAG+ tumor cells were captured using brightfield microscopy. The number of alcian blue positive epithelial cells per field was counted by hand and graphed. The number of mice for each condition is reported as separate dots in graphs. ImageJ was used for quantification of all tumor immunofluorescence images. At least five randomly selected 20x fields of FLAG positive tumor areas were captured for each orthotopic tumor. The average fraction of MUC5AC and GKN2 positive pixels within the mKate2 positive tumor area is reported for the shRNA and TEADi cohorts. For KPC^LOH^ quantification, five random 20x fields of primarily SP cells and five random 20x fields of primarily DP cells were imaged from each mouse. DP regions within SP fields and SP regions within DP fields were manually excluded then the fraction of MUC5AC and GKN2 positive pixels within an mKate2 positive mask area was measured for each field and reported as the average for each mouse.

### Human bulk transcriptomics data analysis

We downloaded publicly available bulk transcriptomics data of 626 surgically resected primary PDAC tumors from 4 studies. RNA-seq data from TCGA (n=142)^74^, CPTAC (n=135)^75^, and ICGC (n=70)^7^ were downloaded from cBioPortal^76^, and microarray data from Puleo et al. (n=279)^6^ were downloaded from ArrayExpress (accession number E-MTAB-6134)^77^. The downloaded gene expression data have already been normalized for library size and log-transformed. We used the *AddModuleScore* function from R package Seurat^73^ to compute expression scores of the gastric, classical, and YAP/TAZ gene programs in each tumor. To account for batch effects between each of the 4 studies, we then normalized the expression scores across tumors in that study such that the normalized scores of each gene program had a mean of zero and equivalent standard deviation. In the heatmap, we further scaled the expression scores across all 626 samples for each gene program such that they range from –1 to 1.

### Mouse KPC^LOH^ bulk RNA-seq data analysis

Partek Flow Software 12.7.0 was used to trim bases with a quality score >20 and minimum read length of 25 bp. STAR 2.7.8a was used for alignment to the mouse genome mm10 and raw gene counts were generated with HTSeq 0.11.0. Count normalization was performed using within-sample normalization by fixing the upper quartile of normalized gene expression counts at 1000 for each sample followed by log2(x+1) transformation. Differential expression analysis was performed using edgeR. To account for differences in baseline expression between animals, matched DP-SP samples from the same mouse were paired to use as a factor variable in the design matrix. A negative binomial generalized linear model (NB-GLM) was fitted using edgeR whereby the response variable was the count data for each gene (unnormalized, unlogged) and the predictor variables were the sample ID and the p53 condition (SP p53 LOH vs DP p53 wild type). We fit the NB-GLM using EstimateDisp and glmQLFit functions and performed differential expression testing by the quasi-likelihood-ratio test using the function glmQLFTest. The differential expression analysis output was used to create a pre-ranked gene list via the log(FDR) with the sign of the fold change for GSEA analysis. For graphing gastric gene enrichment, normalized counts were scaled using a z-score and the mean gastric gene z-score for each SP-DP tumor pair is plotted.

### Analysis of MRTX resistant cell RNA-seq data

For bulk RNA-seq of parental and MRTX1133-resistance human and mouse PDAC cell lines, raw count documents were downloaded from the NCBI Gene Expression Omnibus (accession number GSE269985)^64^. Raw counts were filtered to exclude those with less than 2 maximum counts per sample and normalized using DESeq2 median of ratios. The normalized counts were used for Gene Set Enrichment analysis for each cell line. For RNA-seq of human PDAC lines sensitive or innately resistant to MRTX1133, a pre-ranked gene list was created using the reported differential expression analysis document from supplementary table 4 using the log(FDR) with the sign of the fold change^64^.

### scRNA-seq Analysis of Human PDAC

Human scRNA-seq data was derived from the UCSC Cell Browser PDAC Atlas (https://cells.ucsc.edu/?ds=pdac-atlas) filtering for all epithelial cells across 200 patient samples^78^. YAP/TAZ target genes^68^ and the YAP/TAZ regulated gastric gene signatures were converted to human gene IDs and projected onto UMAPs using the sum of the normalized expression scores.

## SUPPLEMENTARY FIGURE LEGENDS

**Supplementary Figure 1.**
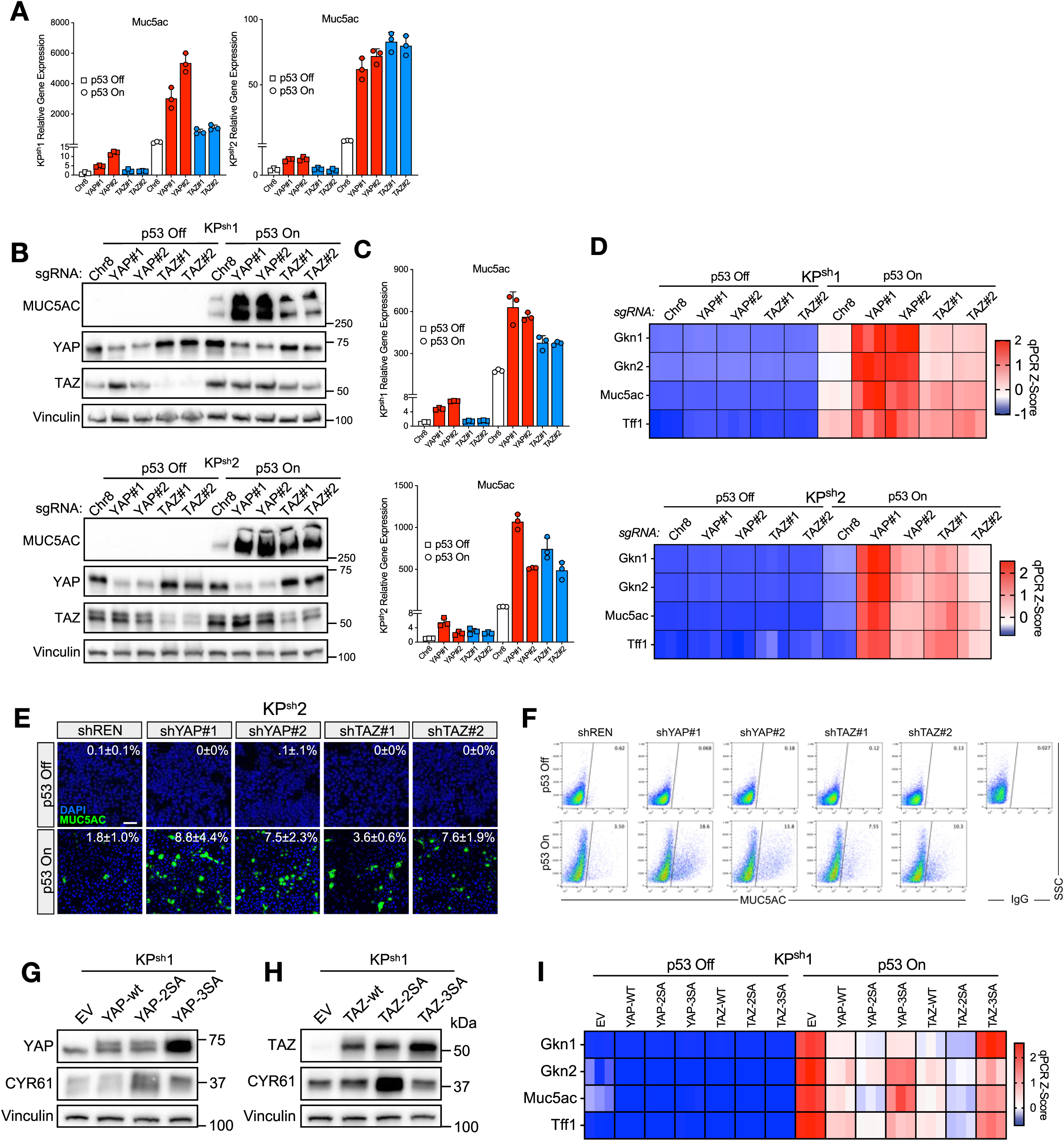
YAP/TAZ activity competes with p53 to control gastric pit-like differentiation. **(A)** RT-qPCRs for Muc5ac in KP^sh^ shYAP, shTAZ, and shREN cells. **(B-D)** Bulk population of KP^sh^ stably expressing sgRNAs targeting Yap1 (YAP), Wwtr1 (TAZ), or an intergenic region of Chromosome 8 (Chr8) and Cas9. **(B)** Representative western blot images showing knockout efficiency and Muc5ac expression. **(C,D)** RT-qPCRs were performed and gene expression is graphed as **(C)** bar graphs for Muc5ac and **(D)** heatmaps of Z-scored expression normalized to Gusb. **(E)** Representative immunofluorescence images of MUC5AC in KP^sh^2 cells. DNA was stained with DAPI. Upper right corner of images displays percentage MUC5AC positive cells with associated standard deviation across an average of n=5.1 fields representing an average of n=2547 cells per condition. **(F)** Scatter plots of representative flow cytometry in KP^sh^1 with indicated shRNAs. **(G-I)** KP^sh^1 overexpressing indicated YAP allele, TAZ allele, or empty vector (EV). **(G,H)** Representative western blots showing **(G)** YAP and **(H)** TAZ overexpression. **(I)** Heatmaps of RT-qPCRs showing Z-scored expression. Scale bar in (E) = 100 µm.

**Supplementary Figure 2.**
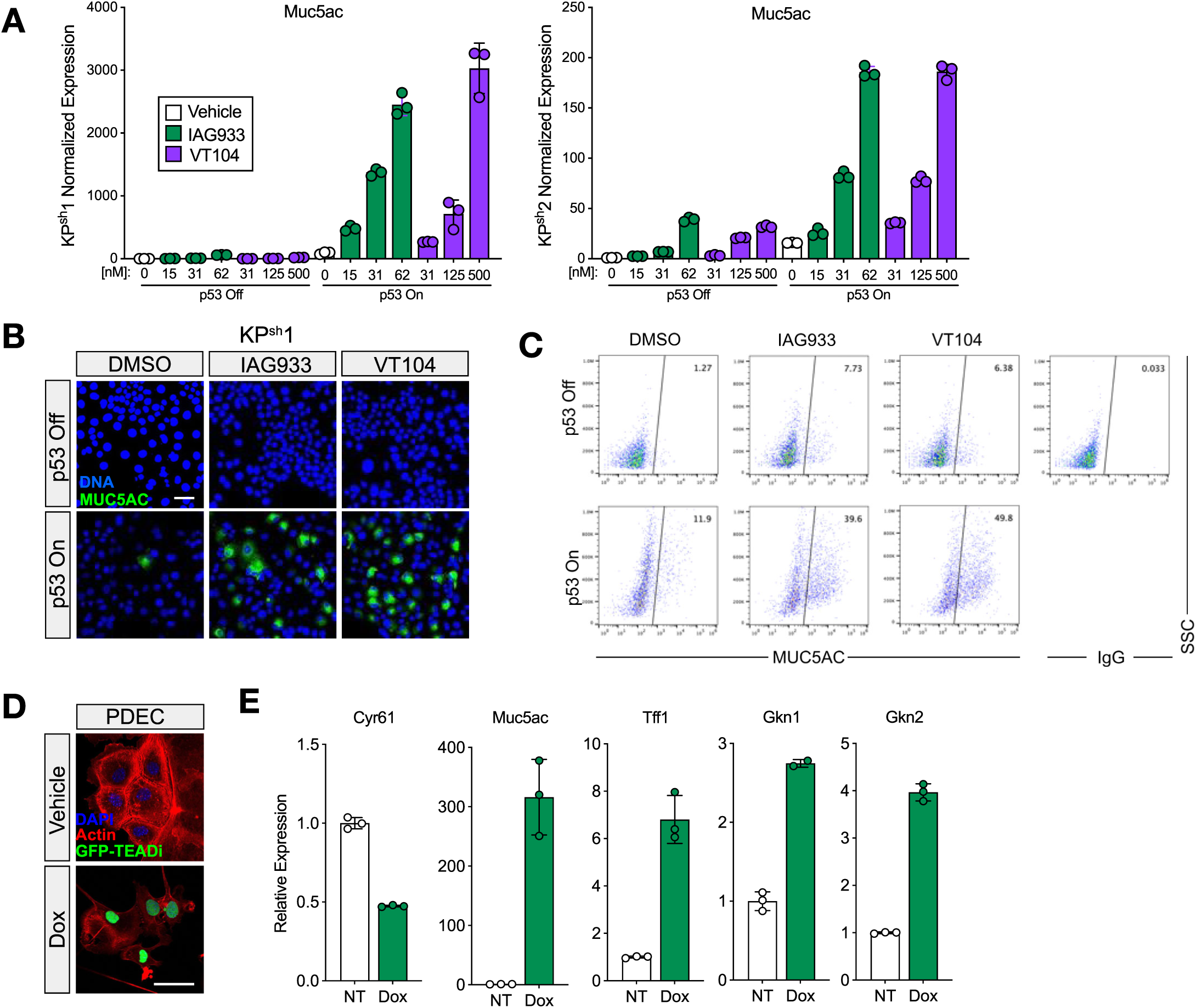
TEAD represses gastric differentiation promoted by p53. **(A**) RT-qPCRs of Muc5ac in KP^sh^ cell lines treated with 48 hours of indicated drugs or DMSO vehicle. **(B)** Representative 40x immunofluorescence images of MUC5AC in KP^sh^1 cells treated with 125 nM IAG933, 500 nM VT104, or DMSO vehicle. DNA was stained with SYTOX Green. **(C)** Representative flow cytometry scatter plots of KP^sh^1 stained with MUC5AC with percentage MUC5AC positive cells in upper right corner. **(D-F)** PDECs stably expressing pInducer20-TEADi-GFP treated with 48 hours of 1 µg/mL dox or vehicle. **(D)** Fluorescence microscopy images of cells stained with Phalloidin (Actin) or DAPI (DNA). **(E)** RT-qPCRs. Scale bar in (B) and (D) = 50 µm.

**Supplementary Figure 3.**
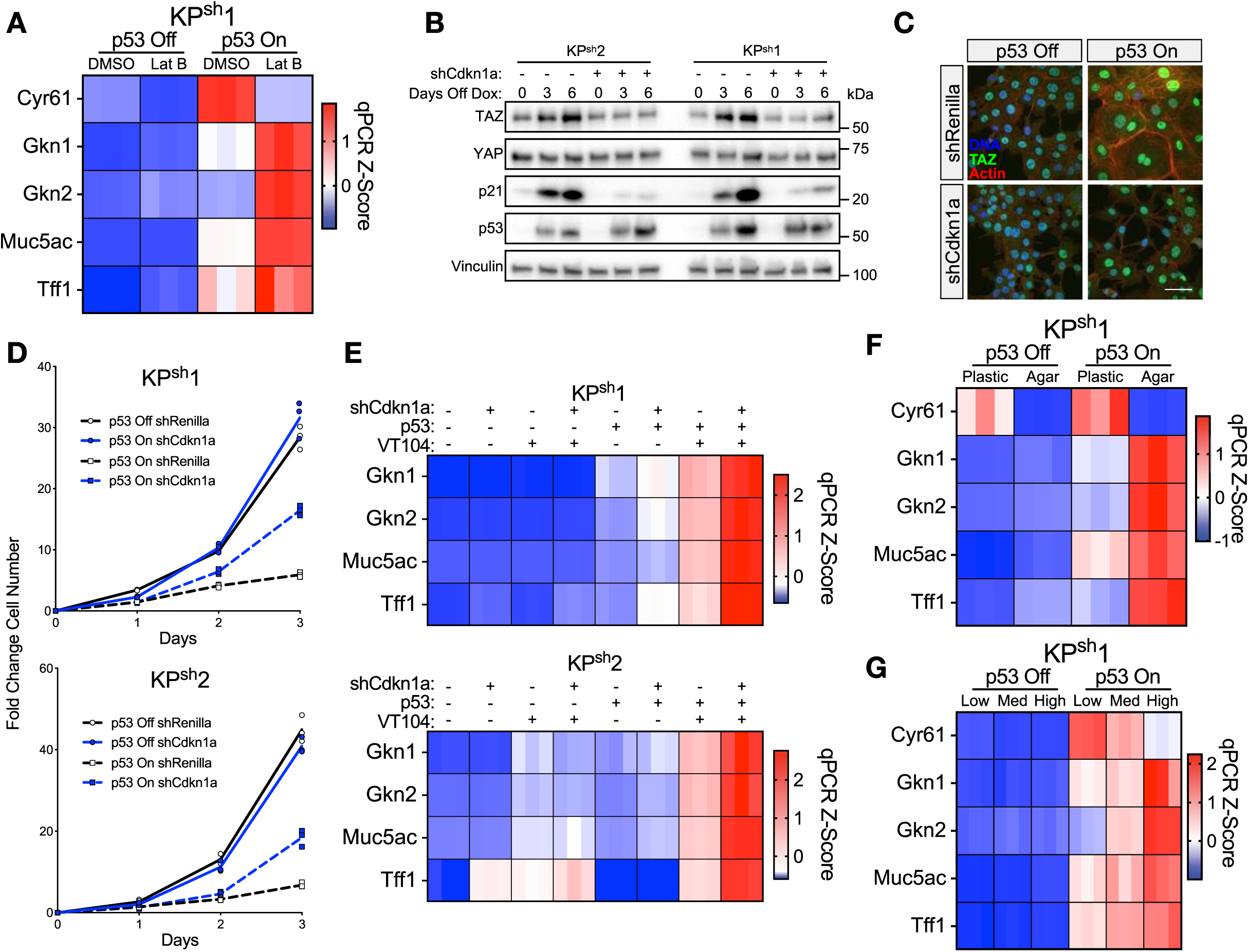
Physiological perturbations of YAP/TAZ activity regulates gastric gene expression. **(A)** KP^sh^1 cells treated with 12 hours of 5 µM Latrunculin B on dox (p53 off) or 6 days off dox (p53 on). Heatmaps of RT-qPCR Z-scored expression normalized to Gusb. **(B-E)** KP^sh^ cells expressing shRNAs targeting Cdkn1a or Renilla luciferase. **(B)** Representative western blots of KP^sh^ cells upon p53 restoration by dox withdrawal. **(C)** Representative IF images on dox (p53 off) or 6 days off dox (p53 on) in KP^sh^2 cells stained with Sytox green (DNA), phalloidin (actin) or a TAZ antibody. **(D)** Growth curves of KP^sh^ cells on dox (p53 off) or following seeding after a 2 day dox washout period. **(E)** Heatmaps of RT-qPCR showing Z-scored expression with 48 hours of 500 nM VT104 treatment or DMS0 control (-). **(F)** KP^sh^1 cells seeded into 0.5% soft agar or on PureCol coated plastic dishes and collected 24 hours after seeding either on dox or at 6 days off dox (p53 on). Heatmaps of RT-qPCR Z-scored expression normalized to Gusb. **(G)** KP^sh^1 cells were seeded at three different cell densities. Heatmaps of RT-qPCR Z-scored expression. Scale bar in (C) = 50 µm.

**Supplementary Figure 4.**
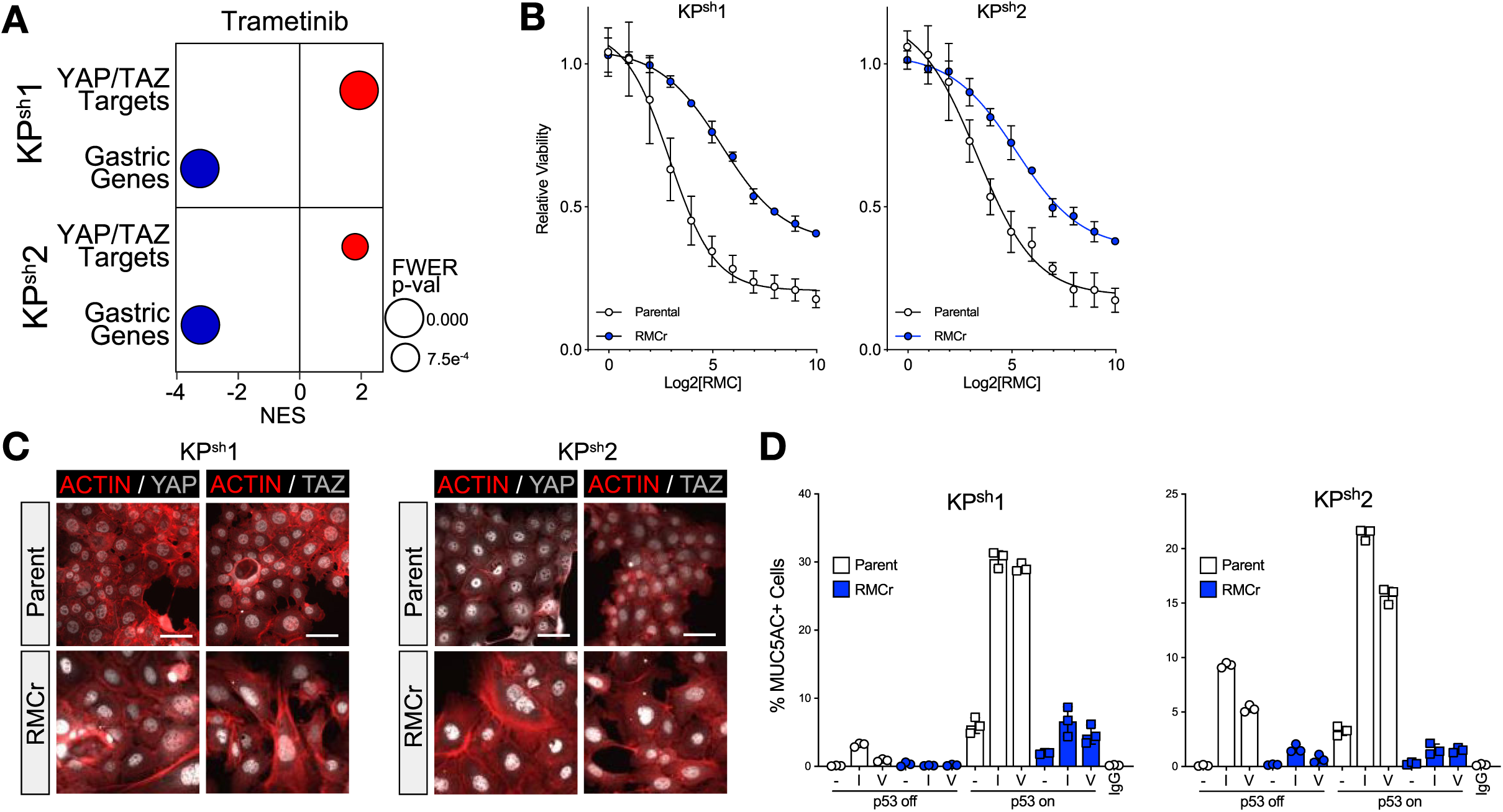
KRAS/MAPK pathway inhibition suppresses gastric gene expression through YAP/TAZ activation. (A) GSEA of RNA-seq in cells treated with 25 nM Trametinib for 72 hours. (B-D) KP^sh^ parental lines or lines resistant to 100 µM RMC-7977 (RMCr). (B) Cell titer glo viability assay in with 96 hours of RMC treatment and (E) IF microscopy images of YAP and TAZ in parental and RMCr KP^sh^ cells. (D) Treatment with 48 hours of 125 nM IAG933, 500 nM VT104, or DMSO showing percentage of MUC5AC positive cells determined by flow cytometry.

**Supplementary Figure 5.**
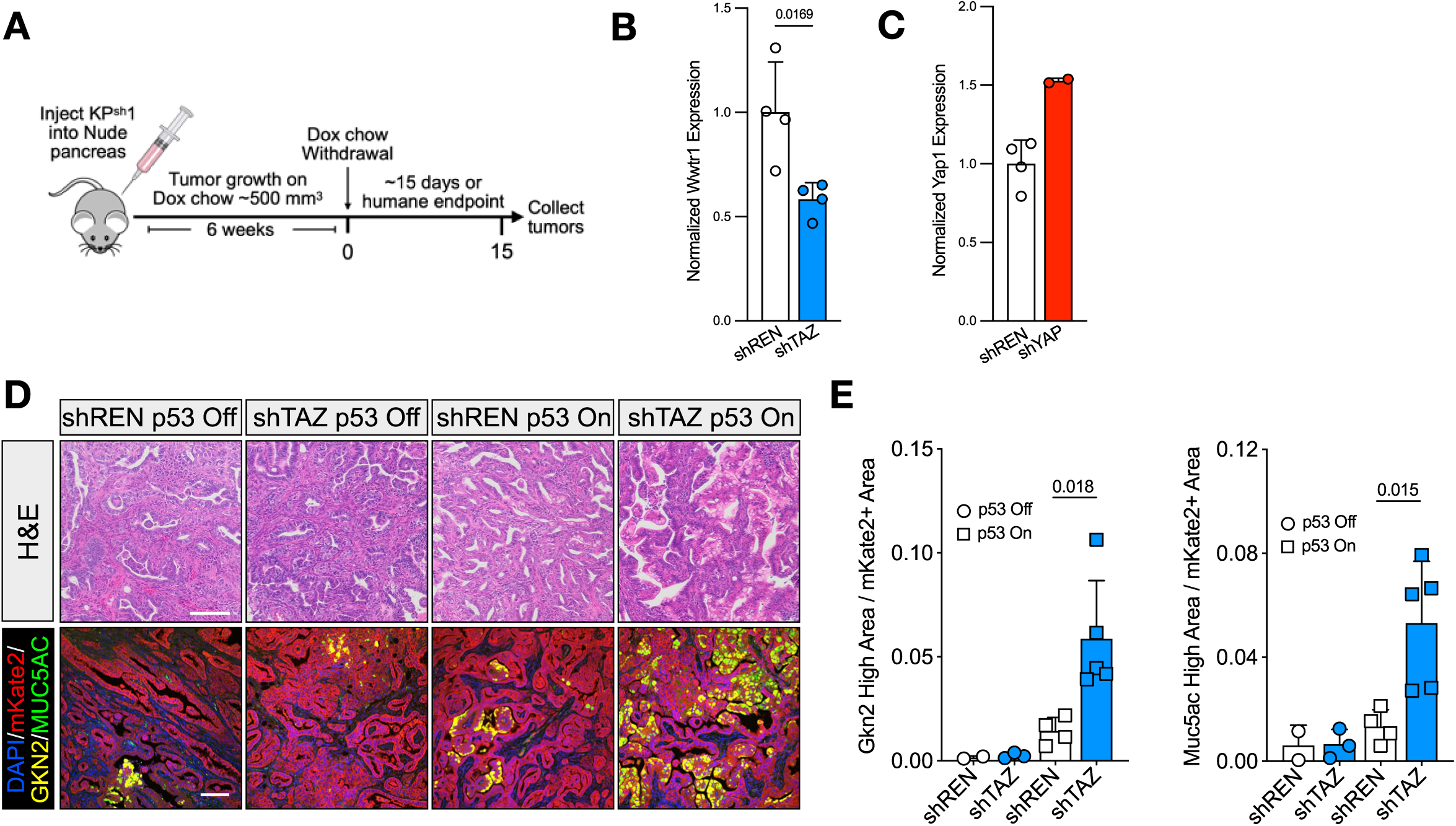
TAZ depletion promotes gastric differentiation in response to p53 restoration in vivo. (A) Schematic of orthotopic pancreas injections of KP^sh^1 cells expressing shRNAs targeting TAZ, YAP, or REN. (B,C) RT-qPCR graphs from mKate2 positive sorted tumor cells following 15 day dox withdrawal demonstrating sustained knockdown efficiency of (B) TAZ shRNA and (C) YAP shRNA. Each dot represents mean expression from technical triplicate wells of a qPCR from cells sorted from one tumor. (D) Representative fields of Hematoxylin and Eosin (H&E) staining (top) or IF microscopy of mKate2, DAPI, GKN2, and MUC5AC (bottom) in shREN vs shTAZ orthotopic tumors on dox (p53 off) or between 9-15 days off dox (p53 on); scale bars = 100 µm. (E) Quantification of GKN2 and MUC5AC positive area per unit mKate2 positive area in shREN and shTAZ orthotopic tumors on dox (p53 off) or between 9-15 days off dox (p53 on). Each dot represents quantification of one tumor from at least five randomly selected fields. P-values calculated by two-tailed students T-test.

**Supplementary Figure 6.**
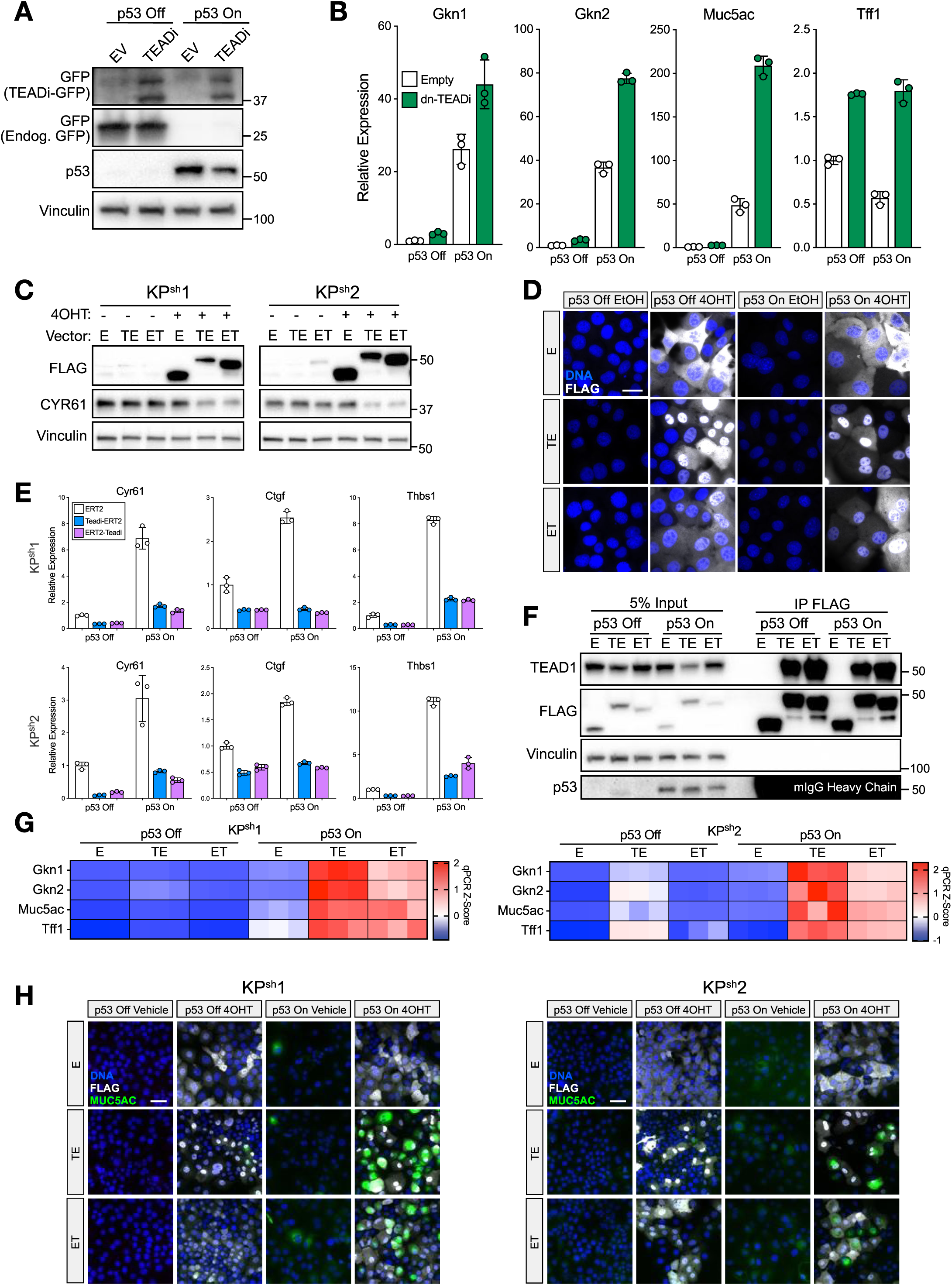
Generation of a tamoxifen inducible dominant negative TEAD. (A,B) TEADi-GFP was constitutively expressed in KP^sh^2 cells and cells were collected. **(A)** Representative western blots showing expression of TEADi-GFP. **(B)** RT-qPCRs of gene expression. **(C-H)** KP^sh^ cells expressing ERT2 (E), TEADi-ERT2 (TE), and ERT2-TEADi (ET) were treated with 48 hours of 1 µM 4-Hydroxytamoxifen (4OHT) or 1:1000 Ethanol vehicle. **(C)** Representative western blots demonstrating 4OHT induction of FLAG linked TEAD inhibitors. **(D)** Representative immunofluorescence microscopy images of KP^sh^1 cells stained with SYTOX green or FLAG antibody; scale bar = 25 µm. **(E)** RT-qPCRs of YAP/TAZ-TEAD target genes. **(F)** Western blots showing FLAG immunoprecipitation (right) or 5% input (left) showing TEAD1 binding to TEAD inhibitor but not control ERT2. **(G)** Heatmaps of RT-qPCRs showing Z-scored expression. **(H)** Representative IF microscopy images demonstrating induction of TEAD inhibition by 4OHT promotes MUC5AC expression; scale bars = 50 µm.

**Supplementary Figure 7:**
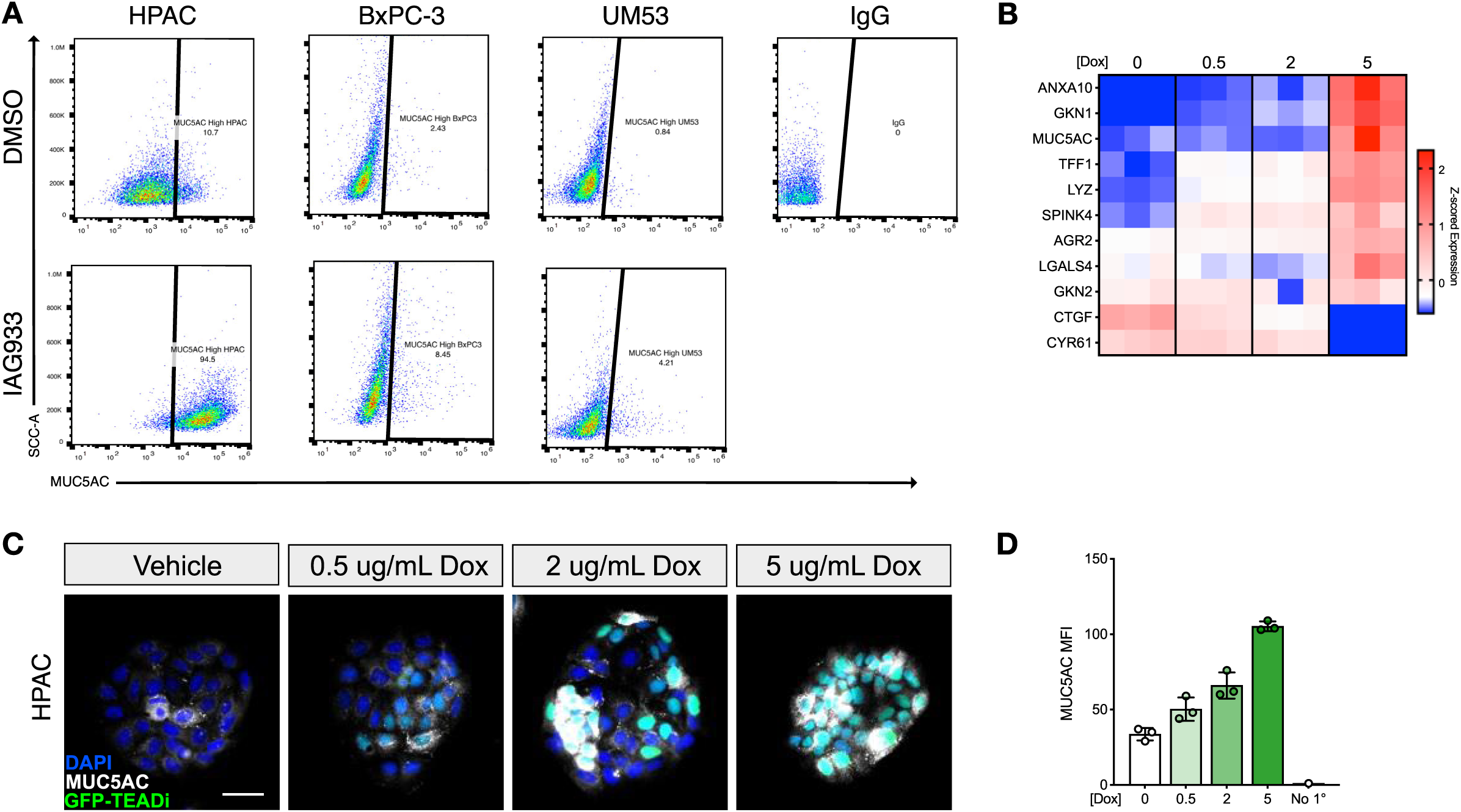
(A) Representative flow cytometry scatter plots of MUC5AC positivity from PDAC cells treated with 1 µM IAG933 for 48 hours. (B-D) HPAC cells expressing pInducer20-TEADi-GFP were treated with indicated µg/mL doses of dox for 48 hours. (B) qRT-qPCRs, (C) representative IF microscopy images, and (D) quantification of flow cytometry median fluorescence intensity of MUC5AC from three-independent wells. Scale bar in (C) = 50 µm

