## Supplementary Table Information for "p53 and YAP/TAZ-TEAD activities determine metaplastic heterogeneity in pancreatic cancer"

Supplementary Table 1 YAP TAZ Target Gene Sets

Supplementary Table 2 Gastric-Like Gene Sets

Supplementary Table 3 Stomach Cell Gene Sets

Supplementary Table 4 shRNA Sequences

Supplementary Table 5 sgRNA Sequences

Supplementary Table 6 dn-TEADi Sequences

Supplementary Table 7 Mus Musculus RT-qPCR Primers

Supplementary Table 8 Human RT-qPCR Primers

Supplementary Table 9 Antibodies
